# The PNUTS-PAD domain recruits MYC to the PNUTS:PP1 phosphatase complex via the oncogenic MYC-MB0 region

**DOI:** 10.1101/2021.12.02.470928

**Authors:** Yong Wei, Alexandra Ahlner, Cornelia Redel, Alexander Lemak, Isak Johansson-Åkhe, Scott Houliston, Tristan M.G. Kenney, Aaliya Tamachi, Vivian Morad, Shili Duan, David W. Andrews, Björn Wallner, Maria Sunnerhagen, Cheryl H. Arrowsmith, Linda Z. Penn

**Affiliations:** Princess Margaret Cancer Centre, University Health Network, 101 College St, Toronto, ON M5G 0A3, Canada; Structural Genomics Consortium, Toronto, Ontario, Canada; Sunnybrook Research Institute, Toronto, ON, Canada; Department of Physics, Chemistry, and Biology, Linköping University, Linköping, Sweden; Department of Medical Biophysics, University of Toronto, 101 College St, Toronto, ON M5G 1L7, Canada

**Keywords:** MYC oncoprotein, PP1:PNUTS phosphatase, direct protein-protein interaction, complex structure, phosphorylation, cancer

## Abstract

Despite MYC dysregulation in most human cancers, strategies to target this potent oncogenic driver remains an urgent unmet need. Recent evidence shows the PP1 phosphatase and its regulatory subunit PNUTS control MYC phosphorylation and stability, however the molecular basis remains unclear. Here we demonstrate that MYC interacts directly with PNUTS through the MYC homology Box 0 (MB0), a highly conserved region recently shown to be important for MYC oncogenic activity. MB0 interacts with PNUTS residues 1-148, a functional unit here termed, <u>P</u>NUTS <u>a</u>mino-terminal <u>d</u>omain (PAD). Using NMR spectroscopy we determined the solution structure of PAD, and characterised its interaction with MYC. Point mutations of residues at the MYC-PNUTS interface significantly weaken their interaction both *in vitro* and *in vivo*. These data demonstrate the MB0 binding pocket of the PAD represents an attractive site for pharmacological disruption of the MYC-PNUTS interaction.

**In Brief:** Solving the structure of MYC-PNUTS direct interaction reveals how the intrinsically disordered MYC-Box0 (MB0) region anchors into a binding pocket in the N-terminal PAD domain of PNUTS. These data provide insight into the molecular mechanism of how the PNUTS:PP1 phosphatase complex regulates MYC phosphorylation.

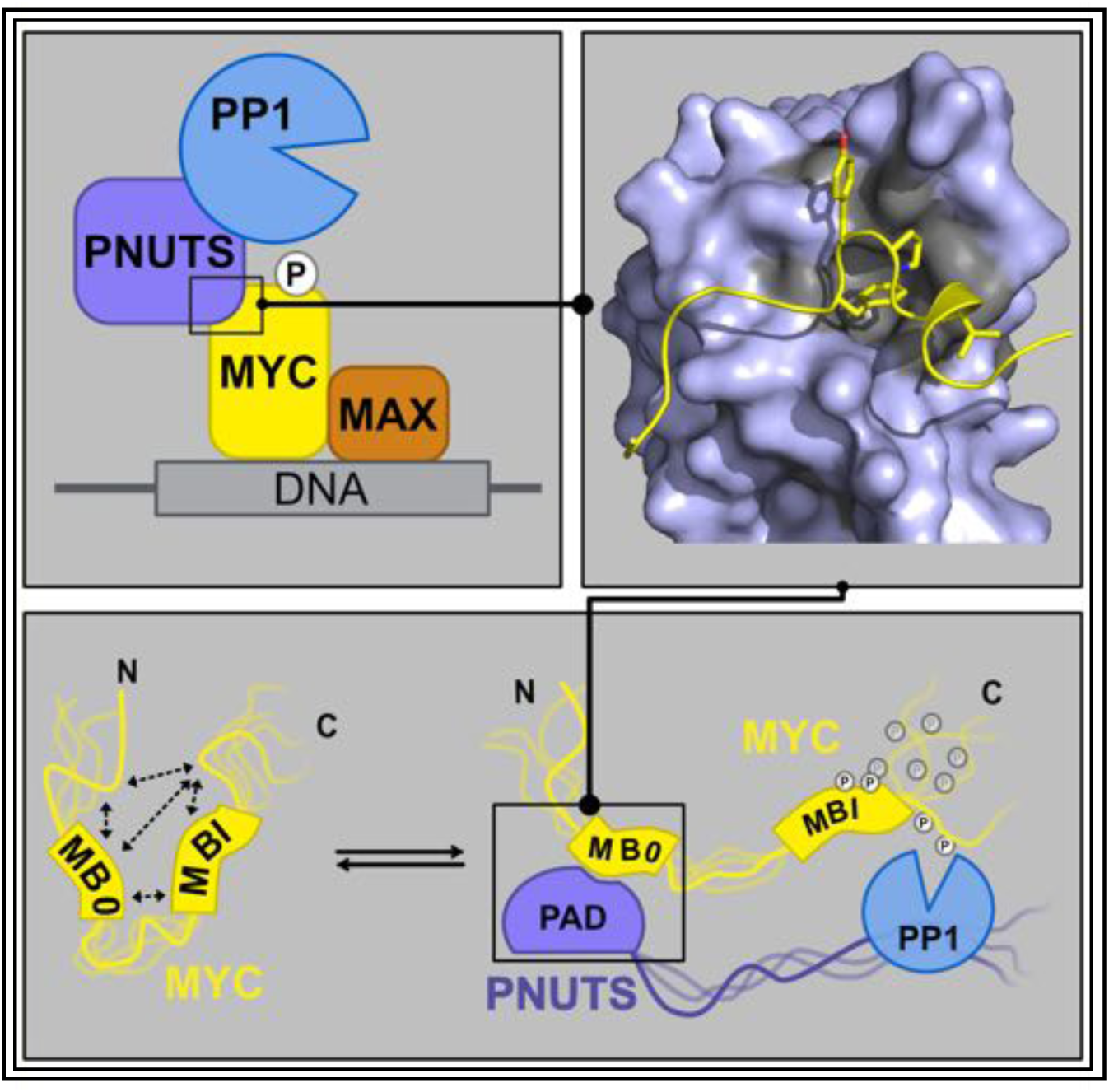

**Highlights:**

- A region critical for MYC oncogenesis, MYC-Box0 (MB0), directly interacts with PNUTS
- <u>P</u>NUTS <u>a</u>mino-terminal <u>d</u>omain (PAD) is a structural domain that interacts with MYC MB0
- Mutation of single residues at the interaction interface disrupts MYC-PNUTS binding in cells
- MYC-PNUTS binding releases MYC intramolecular interactions to enable PP1substrate access

## Introduction

Dysregulated MYC activity is a hallmark of >50% of human cancers and is often linked to aggressive disease and poor prognosis(Kalkat et al., 2017; Meyer and Penn, 2008). MYC controls the transcription of ∼15% of genes, thereby regulating numerous biological processes including cell growth, metabolism and immune response(Casey et al., 2017; Lorenzin et al., 2016; Luscher and Vervoorts, 2012; Meyer and Penn, 2008; Poole and van Riggelen, 2017). In non-transformed cells, the expression of this master-regulator is tightly controlled. By contrast, in cancer MYC activity is dysregulated by a plethora of mechanisms, resulting in constitutive activity that drives oncogenic growth. The MYC family of proteins also includes N-MYC and L-MYC, whose expression is normally restricted to fetal development, but can be reactivated and dysregulated in cancer.(Meyer and Penn, 2008) Thus, dysregulated MYC activity is a potent oncogenic driver of most human cancers.

Evidence from mouse models of cancer strongly suggest that inhibiting MYC oncogenic activity will dramatically improve cancer outcome(Ponzielli et al., 2005; Prochownik and Vogt, 2010; Whitfield et al., 2017), as treatment with cycles of systemic genetic suppression of MYC leads to tumour eradication, without adverse side-effects to normal cells(Bellovin et al., 2013; Felsher and Bishop, 1999; Li et al., 2014; Soucek et al., 2013). Despite these promising results, targeting MYC using traditional drug development approaches has failed(Chen et al., 2018; Dang et al., 2017; McKeown and Bradner, 2014), largely because MYC is intrinsically disordered in the absence of a binding partner(Tu et al., 2015). MYC contains a basic helix-loop-helix leucine zipper domain (bHLHLZ), common to many transcription factors, and six regions termed MYC boxes that are unique and highly conserved amongst the MYC family of proteins. MYC Box II (MBII), and more recently MYC Box 0 (MB0), have been shown to be functionally required for the full oncogenic activity of MYC, suggesting these MYC boxes are key regulatory regions(Kalkat et al., 2018). A promising strategy to develop MYC inhibitors is to identify MYC protein interactors that are essential for MYC oncogenic activity, and then disrupt these MYC-protein interactions by targeting the structured region of the partner proteins(Bugge et al., 2020; Lu et al., 2020). The identification of structurally and dynamically unique binding modes of MYC to different protein interactors will then lead to the development of an arsenal of inhibitors targeting MYC activity with high specificity. However, few direct protein interactors of MYC have been validated. As a first step to filling this gap, we identified hundreds of novel MYC binding proteins using the BioID in-cell, proximity labelling technique followed by mass spectrometry, which includes both direct and indirect protein interactors(Dingar et al., 2015; Kalkat et al., 2018). We then further characterized one of the hits, protein phosphatase 1 (PP1), a serine threonine phosphatase and its regulatory substrate-specifying subunit, PP1 nuclear targeting subunit (PNUTS), as we and others have shown that phosphorylation regulates MYC stability and/or activity(Huang et al., 2004; Wasylishen et al., 2013)(20-23)(Chakraborty et al., 2015; Hemann et al., 2005). Indeed, inhibiting PP1 using RNAi or pharmacological inhibitors triggers MYC hyperphosphorylation, leading to chromatin eviction and MYC protein degradation(Dingar et al., 2018). Exploiting this PP1:PNUTS-MYC regulatory axis to target MYC destruction has promise; however, inhibiting PP1 catalytic activity is not a viable approach as PP1 has several protein substrates(Bollen et al., 2010; Casamayor and Arino, 2020; Cohen, 2002). Thus, to advance our understanding of how MYC is regulated by this phosphatase complex and to evaluate the potential for pharmacological disruption of MYC interaction with PP1:PNUTS to promote MYC degradation, we sought to delineate the molecular basis of this interaction.

Here, we report that MYC interacts directly with PNUTS. Using biolayer interferometry (BLI) analysis the interaction has been mapped to conserved residues 16-33 of MYC, recently termed MB0(Helander et al., 2015), and a functional unit of PNUTS consisting of the N-terminal 148 residues, which we have termed the <u>P</u>NUTS <u>a</u>mino-terminal <u>d</u>omain (PAD). Using nuclear magnetic resonance (NMR) spectroscopy, we first determined the structure of PAD alone (apo). Building on these results, the molecular interaction of PAD and MB0 was then resolved by analyzing each protein in the presence of the partner protein, and by analyzing a PAD-MB0 fusion protein, in conjunction with molecular modeling. Together, this revealed the critical residues and structural details essential for the PNUTS-MYC interaction. Validation of key residues important for the interaction within both proteins was achieved by demonstrating that point mutants of these residues disrupted the PNUTS-MYC interaction *in vitro* and *in vivo*. Taken together, these results not only provide new insights into the molecular basis of MYC interaction with PNUTS, but also provide foundational data for the potential development of drugs targeting PNUTS to disrupt the PNUTS-MYC interaction and inhibit MYC oncogenic activity.

## Results

### MB0 interacts directly with PAD

Our previous work identified MYC as a substrate of the PP1:PNUTS phosphatase complex, however the molecular basis of the interaction and whether it is direct remained unclear(Dingar et al., 2018). As MYC does not contain a canonical PP1 recognition motif (RVxF), we hypothesized that MYC may interact with the non-catalytic, substrate-specifying subunit PNUTS. PNUTS contains two previously annotated domains: a TFIIS helical bundle-like domain (residues 73-147) of the Med26 Pfam family (PF08711) with no functional annotation(Zacharchenko et al., 2016); and a zinc finger domain at the C-terminus(Allen et al., 1998; Kim et al., 2003) (Fig. 1a). Since the former is a putative protein-protein interaction domain, we first interrogated the interaction potential of the N-terminal region by designing and testing several expression constructs of proteins within the first 186 amino acids of PNUTS. The shortest of these that was successfully expressed and purified as a stable protein encompassed residues 1-148 (PNUTS(1-148)), suggesting this region represents an independently folded functional domain. This N-terminal region of PNUTS(1-148) was then further evaluated in binding assays with MYC peptides and protein.

**Figure 1.**
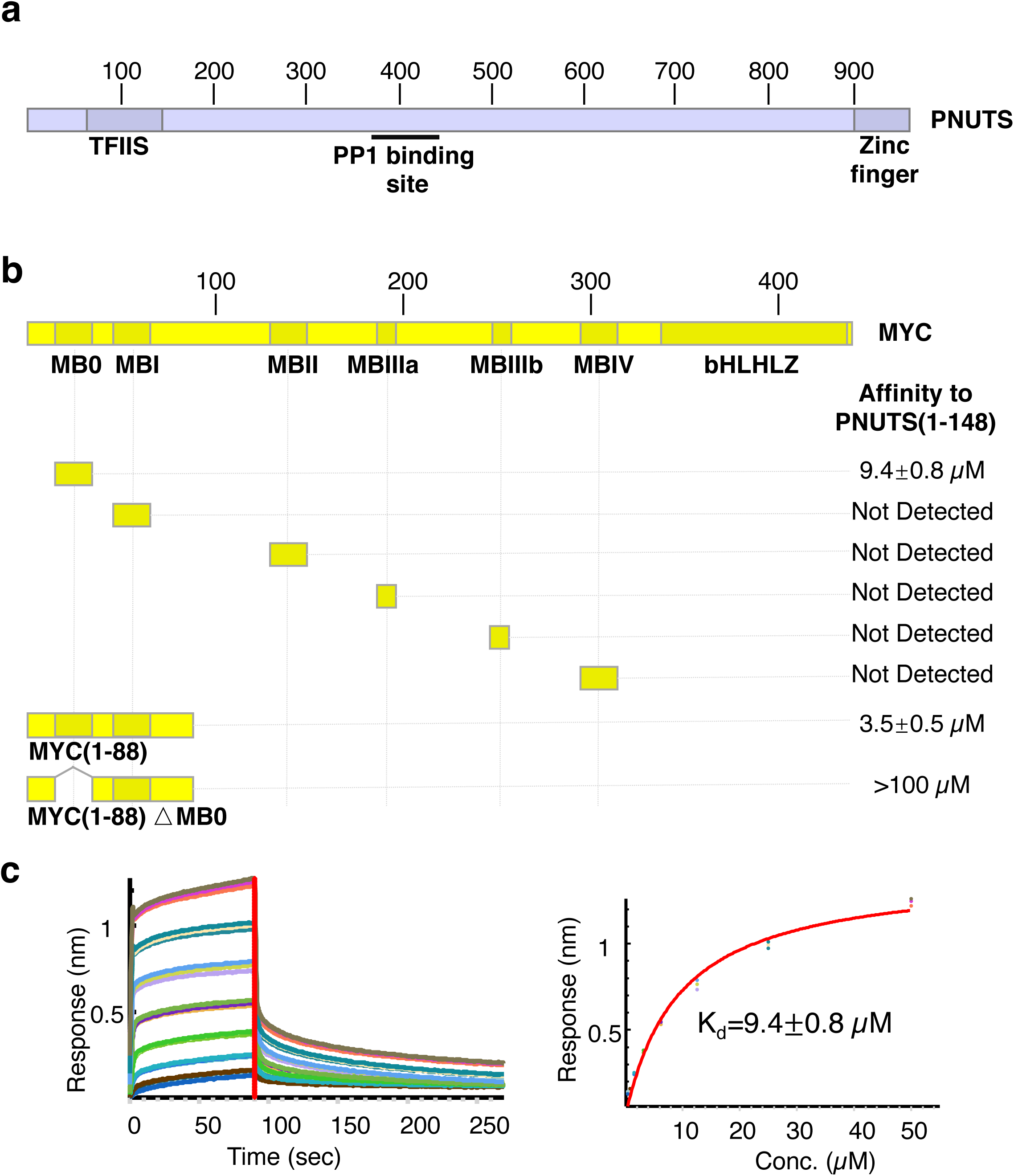
MB0 interacts directly with PAD. (**a**) Domain organization of PNUTS. Two previously annotated domains (TFIIS and Zinc finger) are highlighted with blue boxes and the PP1 binding region is indicated. (**b**) Domain organization of MYC with homology boxes (MB0, MBI, MBII, MBIIIa, MBIIIb, MBIV) as well as the basic helix-loop-helix leucine zipper domain (bHLHLZ) highlighted in dark yellow (**top**). The different MYC constructs that were tested by BLI for binding to PNUTS(1-148) are displayed below with the measured affinities denoted on the right. **(c)** Representative BLI sensorgrams (**left**) and steady-state curve fit (**right**) for MB0 binding to PNUTS(1-148). See also Figure S1.

To determine whether PNUTS(1-148) interacted directly with MYC, we initially focused our analysis on the six highly conserved and regulatory MYC Box (MB) regions. BLI was used to evaluate the binding of each MB (MB0, MBI, MBII, MBIIIa, MBIIIb, MBIV) peptide to PNUTS(1-148). Briefly, biotinylated MB peptides were immobilized onto streptavidin-conjugated biosensors and subsequently suspended in a serially diluted PNUTS(1-148) protein solution. The BLI signals were then used to calculate apparent binding affinities. MB0 interacted with PNUTS(1-148) with a K_d_ of 9.4 ± 0.8µM, whereas the other MYC boxes failed to produce detectable signals (Fig. 1b; Supplementary Fig. 1a-e), suggesting that MB0 constitutes the primary interaction site of MYC with PNUTS(1-148). We next performed reciprocal BLI assays to test the interaction of recombinant biotinylated MYC N-terminal protein, comprising residues 1-88 (MYC(1-88)) that contains MB0 and MBI, and its deletion mutant lacking MB0 (MYC(1-88ΔMB0)). MYC(1-88) directly interacted with PNUTS(1-148) with a K_d_ of 3.5±0.5 µM, while MYC(1-88ΔMB0) demonstrated a weak but still measurable interaction with PNUTS(1-148) (K_d_ >100 µM) (Fig. 1b; Supplementary Fig. 1f, g). These data suggest that while regions outside of MB0 can contribute to binding, MB0 is the dominant PNUTS anchor site. Having established the direct interaction of MB0 with PNUTS(1-148), and since the folding of this entity requires sequence components beyond the PNUTS TFIIS-like sequence, we termed this newly identified region the <u>P</u>NUTS <u>a</u>mino-terminal <u>d</u>omain (PAD).

### Specific residues within MB0 interact with PAD

To identify MB0 residues critically involved in binding to PAD, we used NMR spectroscopy. We have previously shown that MB0 and MBI are transiently ordered regions, in an otherwise disordered 6His-MYC(1-88) fragment, which was refolded from inclusion bodies(Andresen et al., 2012). For this study, we expressed and purified MYC(1-88) under native conditions from a cleavable TRX-tagged construct, and found the NMR chemical shifts of ^13^C-^15^N-MYC(1-88) near-identical to those of 6His-MYC(1-88). NMR titrations were then performed with unlabeled PAD at 15°C. As MYC(1-88) is predominantly intrinsically disordered, most of its amide resonances in the (^1^H-^15^N) HSQC spectra overlap heavily and are clustered between 7.7 and 8.6 ppm(Andresen et al., 2012; Fladvad et al., 2005). Nevertheless, an (^1^H-^15^N) HSQC overlay at increasing PAD:MYC(1-88) ratios shows clear shifting and/or broadening of V19, Q20 and F23 peaks (Fig. 2a).

**Figure 2.**
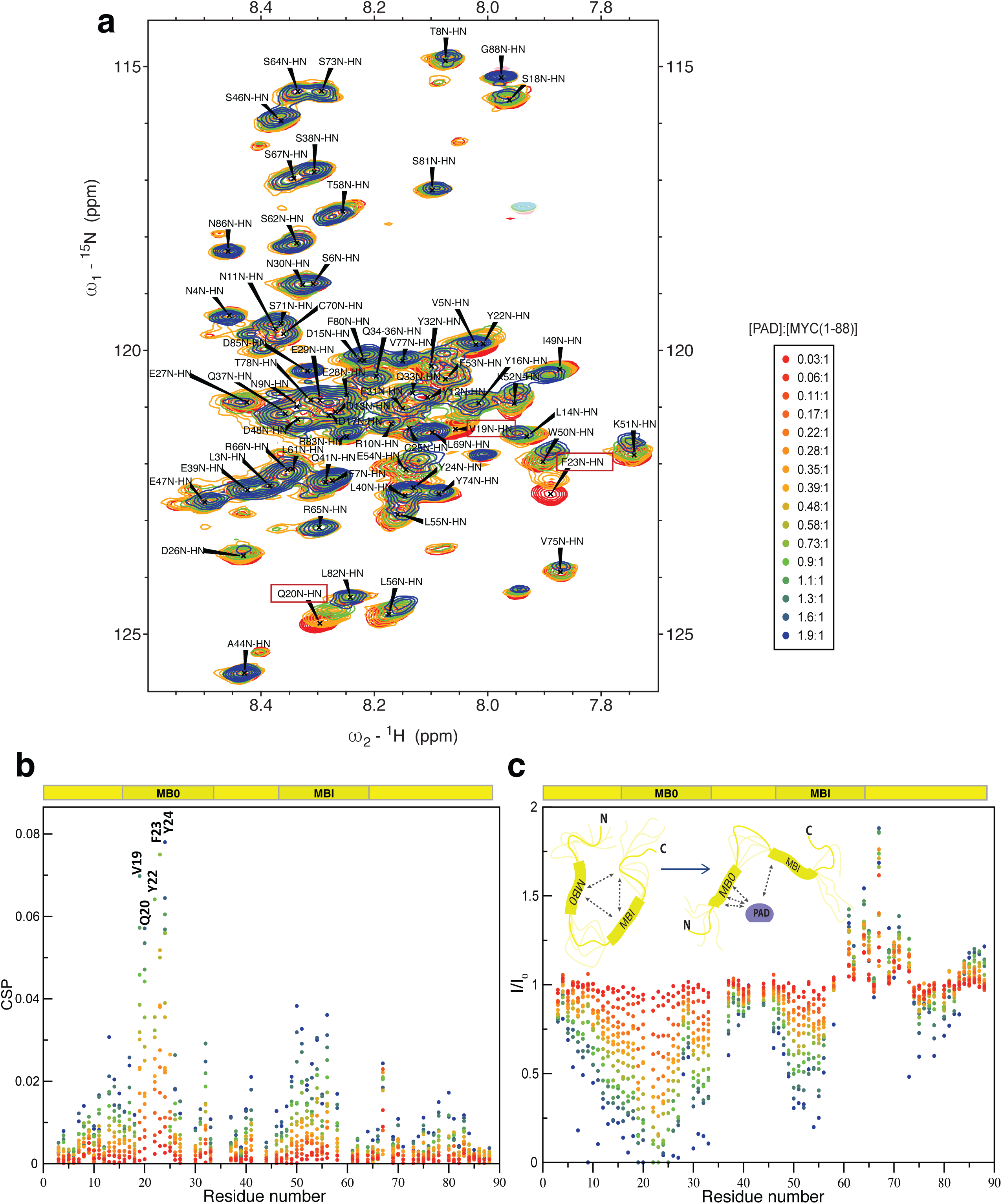
Specific residues within MB0 interact with PAD. (**a**) ^1^H-^15^N HSQC spectral overlay of MYC(1-88) in the presence of increasing concentrations of PAD (left). The color legend for panels a-c is displayed (right). (**b**) Composite CSPs of MYC amide resonances from HNCO spectra upon addition of PAD at MYC-to-PAD molar ratios ranging from 0.06 to 1.9. MYC. Residues with significant CSPs are labeled, with MB0 and MBI regions in MYC(1-88) highlighted by dark yellow boxes below the graph. (**c**) Changes in intensity ratios (I/I0) of MYC(1-88) amide resonances from same HNCO spectra as in (b), and with MB0 and MBI regions similarly highlighted below the graph. Cartoon inset illustrates how increased I/I_0_ for residues in and adjacent to MBI, reflect more rapid interconversion between states in this region than in free MYC(1-88), agreeing with PAD-induced release of transient interactions between MB0 and MBI regions present in the unbound state. See also Figure S2.

To more accurately determine chemical shift perturbations (CSPs) and broadening of MYC amide resonance due to PAD binding, it was necessary to acquire conventional 3D backbone spectra (HNCO and HNCA) to resolve the amide signals based on their coupling to neighbouring carbonyl groups or Cα carbons. When MYC(1-88) was titrated with PAD, substantial CSPs and broadening of MYC resonances were observed (Fig. 2b, c). A six-residue stretch of MB0 (V19 to Y24) exhibited the most substantial CSPs and broadening effects (Fig. 2b), which is in agreement with MB0 being critical for PAD binding by BLI (Fig. 1b). All assigned MYC residues between D13 and Q33 lose at least 75% of their peak intensity at a 1.9:1 PAD to MYC ratio with no signs of signal recovery in the bound state (Fig. 2c, Supplementary Fig. 2). This indicates continued chemical exchange between multiple bound states(Bozoky et al., 2013; Helander et al., 2015), which is characteristic for intrinsically disordered proteins(Forman-Kay and Mittag, 2013; Fuxreiter, 2018). Consistent with our BLI data, other parts of MYC(1-88) also show resonance shifts and broadening, including a sequence in MBI (residues W50-L55) with smaller CSP and intensity changes compared to those of MB0. Interestingly, a region within and adjacent to MBI (L61 to V75) showed sharper amide signals in response to PAD binding, with S64, R65 and S67 amide intensity increased up to 6-fold (Fig. 2c). This suggests that PAD interaction with MB0 leads to greater conformational mobility of this segment and more rapid interconversion between states than in free MYC(1-88). This would be in agreement with PAD-induced release of previously identified transient interactions between MB0 and MBI regions (Andresen, 2012) (Fig. 2c, inset).

### PAD adopts a helical bundle with an Armadillo subdomain

To understand how PNUTS recognizes and binds to MYC, we first determined an NMR-derived solution structure of PAD (PDB ID: 6VTI, Table 1; Supplementary Fig. 3). The topology of the overall structure consists of nine α-helices (α1, aa8–18; α2, aa28–39; α3, aa44–56; α4, aa60–69; α5, aa71–85; α6, aa88–100; α7, aa105–111; α8, aa113-123; α9, aa127-147; Fig. 3a, b) that form a series of helical bundles. These can be grouped into two subdomains: i) an <u>N</u>-terminal <u>d</u>omain (NTD), consisting of α1-3; and, ii) an Armadillo (ARM) subdomain (aa60-148). This ARM subdomain is composed of two consecutive ARM repeats (ARM1 and ARM2), made up of α4-6, and α7-9, respectively (Fig. 3b). ARM repeats, commonly associated with protein interactions (Pfam family PF00514) are made up of three α-helices including a short α-helix (for example, α7 of ARM2) and two longer α-helices (α8 and α9 of ARM2) (Fig. 3b).

**Figure 3.**
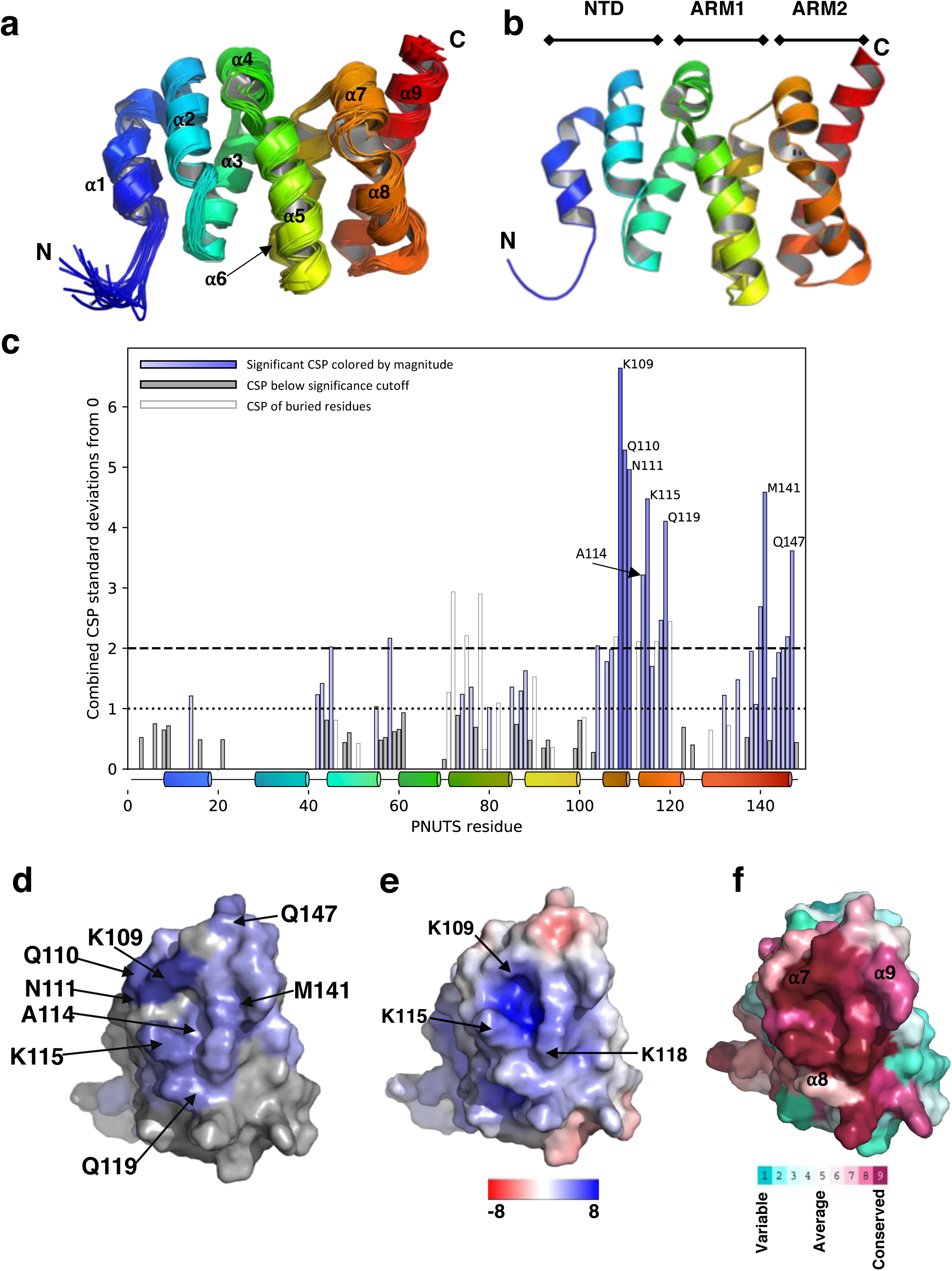
MB0 interacts with the C-terminal facing surface of PAD ARM2 motif. (**a**) Backbone trace of the NMR ensemble of PAD (20 structures), as seen from the ‘front’ view. Helices are numbered with respect to their order from the N- to C-terminus colored from blue to red. (**b**) Cartoon representation of the lowest energy PAD structure, which comprises an N-terminal terminal domain (NTD) consisting of a 3-helix bundle (helices 1 to 3), followed by two Armadillo (ARM) repeats (ARM1 and ARM2, helices 4 to 6 and 7 to 9, respectively). (**c**) Combined chemical shift perturbations δ_i_ for each residue i of PAD upon complex formation with MB0 peptide and MYC(1-88) protein. The δ are normalized to 1 standard deviation from 0, see Methods under Molecular Docking of MB0 to the PAD. Residues with significant δ_i_, differing by more than 1 standard deviation from 0, are colored blue, with a more saturated hue for greater δ_i_. Residues with a δ_i_ lower than 1 standard deviation from 0 are colored grey. Bars of buried residues are colored white with grey borders. A residue is defined as buried if less than 5% of its relative area normalized as a Gly-X-Gly chain is exposed. (**d**) Surface representation of apo-state PAD as seen from the view of a 70 degree rotation of panel (a) on the Y axis, with residues colored by the scheme used in Figure 3c. (**e**) Representation of the PAD electrostatic surface potential using a color gradient spanning red (kT/e =-8) to blue (kT/e =8), in the same view as (**d**). (**f**) Analysis of PAD sequence conservation between species (using ConSurf server) with a structure color-coding bar spanning turquoise (for residues that are not conserved) to maroon (highly conserved residues), in the same view as (**d**). See also Figure S3 and S4.

**Table 1.**
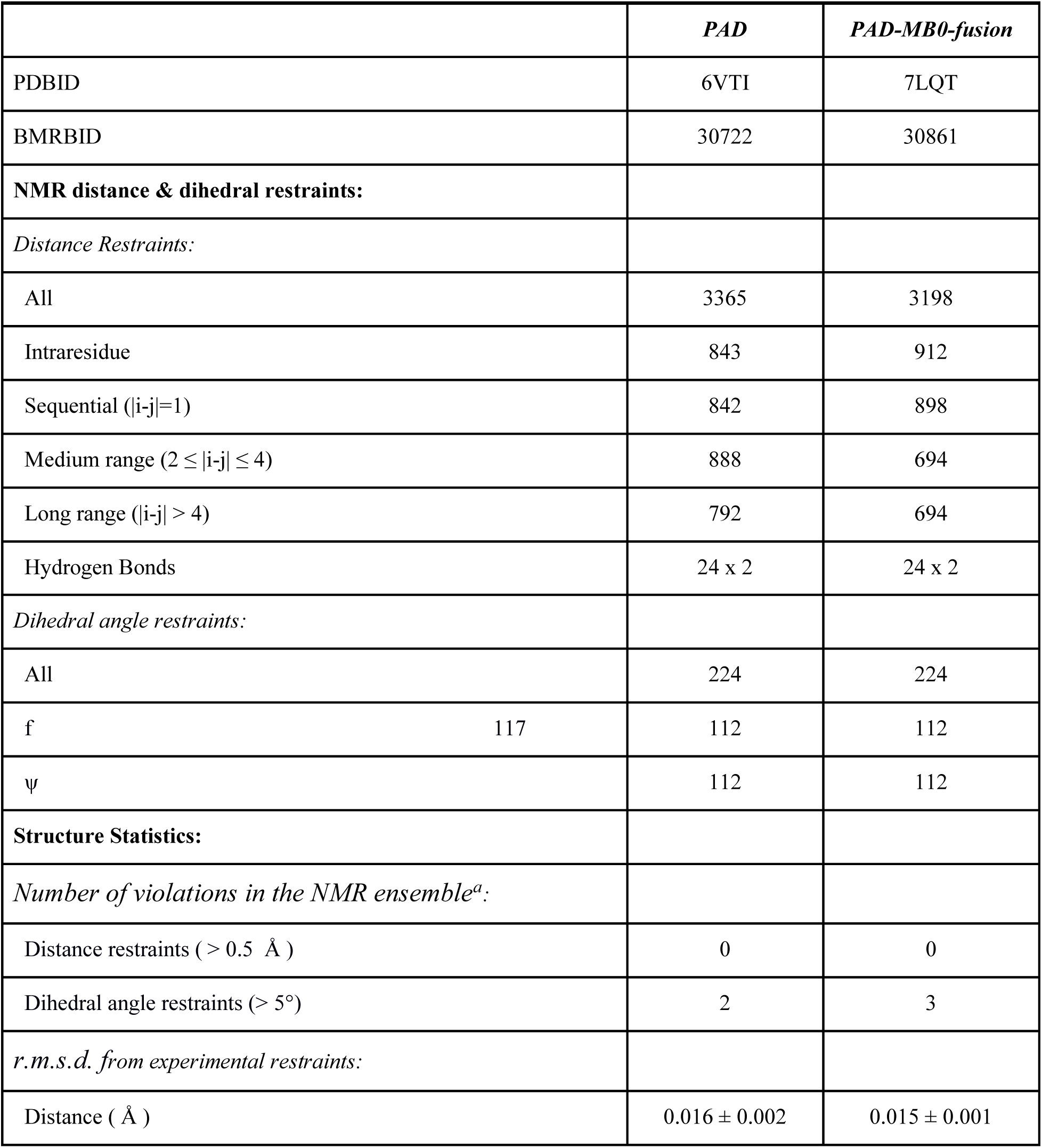
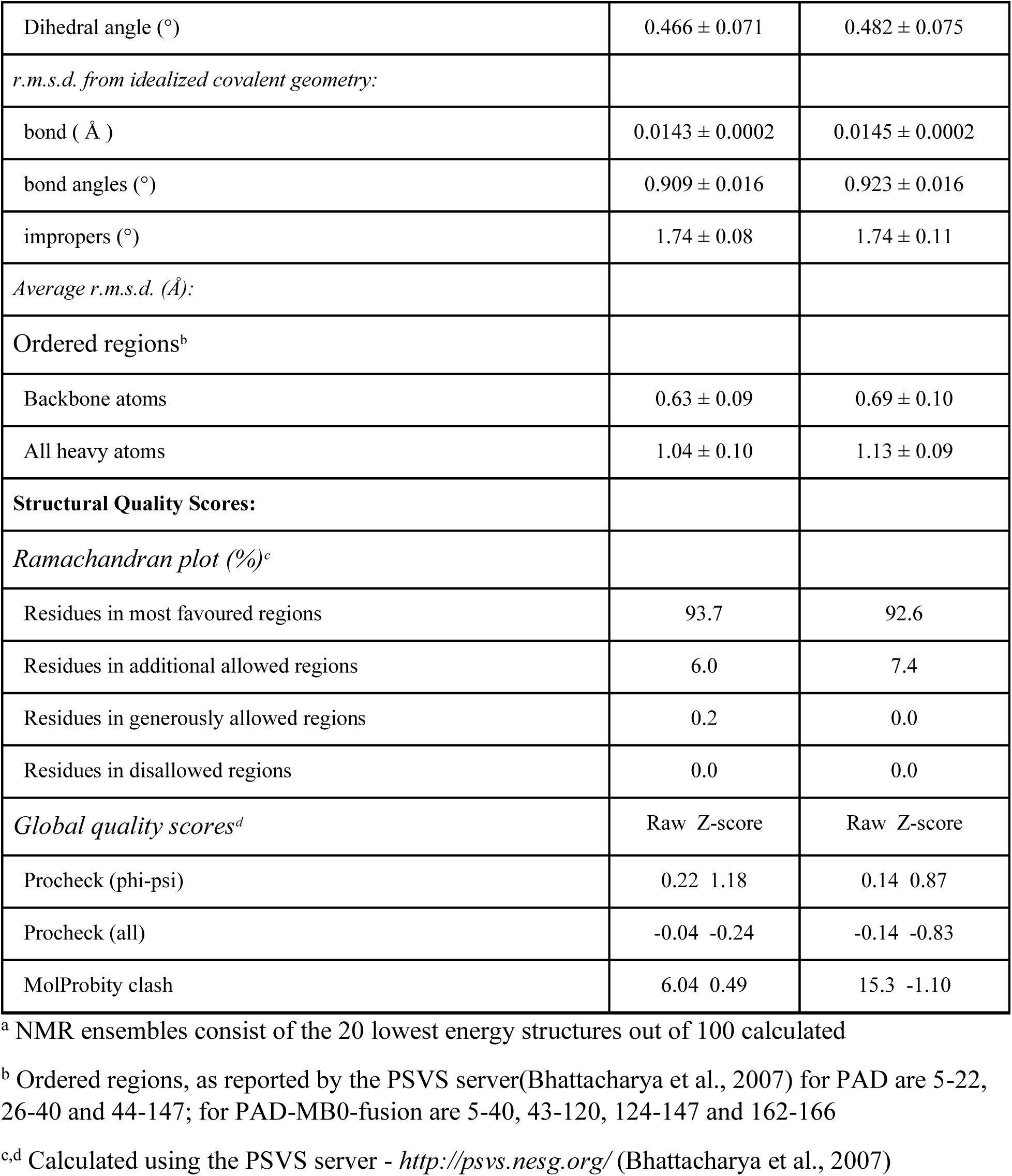
Statistics of NMR constraints and structural parameters for the NMR ensembles of apo PAD and PAD-MB0-fusion complex.

### MB0 interacts with the C-terminal facing surface of PAD ARM2 motif

To map the region of interaction between MB0 with PAD, we performed NMR binding studies at 30°C using ^13^C, ^15^N-labeled PAD titrated with biotinyl-MB0 peptide and MYC(1-88) (Supplementary Fig. 4a, b). At a MB0 to PAD molar ratio of 2 to 1, there was a significant movement of PAD amide resonances in (^1^H-^15^N) HSQC spectra with an average CSP of 0.012 ppm and a STDEV of 0.018. A similar perturbation pattern was observed, when MYC(1-88) was added to PAD (Supplementary Fig. 4b). We then analyzed normalized combined CSPs for these experiments (Fig. 3c, see Methods for CSP calculations). Eight residues (K109, Q110, N111, A114, K115, Q119, M141, and Q147) had composite amide CSPs ≥ 3 standard deviations from 0, and an additional 13 had values ≥ 2 standard deviations from 0, of which 6 are solvent-exposed according to the NMR-derived solution structure (V45, R58, T104, K118, W140, and S146) (Fig. 3c). These data further confirm that the MB0 residues are the dominant drivers of the PNUTS-MYC interaction, consistent with our BLI data (Fig. 1b and 1c) and NMR titrations of (^13^C, ^15^N) MYC(1-88) (Fig. 2). A map of CSPs on the PAD surface in response to MB0 interaction indicates that the highly perturbed residues (with a combined CSP greater than 1 standard deviation) form part of, or are close to the C-terminal facing surface of the ARM2 motif (Fig. 3d). This C-terminal facing surface is mainly composed of hydrophobic residues in the center, and surrounded by several charged residues, including three Lysines (K109, K115, and K118) (Fig. 3e; Supplementary Fig. 4c). We next used (^1^H-^13^C) HSQC spectra of the PAD, to detect perturbations to the resonances of amino acid side chains. The methyl groups of A114 (γ), I144 (δ1 and γ2), T113 (γ2), and M141 (ε) – all in or near the C-terminal facing surface - were noticeably perturbed upon MB0 peptide binding (Supplementary Fig. 4d). Importantly, a ConSurf analysis(Ashkenazy et al., 2016; Landau et al., 2005) indicates that the C-terminal facing surface of the ARM2 motif is conserved across species (Fig. 3f), suggesting a potential conserved interaction surface.

### NMR guided model of PAD and MB0 interaction

To analyze structural features of the PAD-MB0 interaction, a computational modelling approach was taken to identify possible interaction modes of MYC with PNUTS. Guided by the intensity changes and CSPs of MB0, and the CSPs of PAD as described above (see methods), we generated 50,000 models of the PAD-MB0 complex using the Rosetta FlexPepDock ab-initio protocol(Raveh et al., 2011), with NMR-derived constraints accounting for circa 50% of the inter-chain interaction energy. These models were clustered with respect to the position of MB0 residues 19-26 and, using a LASSO algorithm(Tibshirani, 1996) the combination of the fewest possible clusters which together best satisfy the experimental constraints was determined. Nine clusters are required to describe 71% of the variance evident in the experimental data (Supplementary Fig. 5a, https://modelarchive.org/doi/10.5452/ma-3ef73 code: LWSnMfLXwf), with one dominant cluster (Cluster 0) representing 28.5% of all models (Supplementary Fig. 5b,c).

The resulting NMR-driven ensemble of PAD-MB0 complexes (Fig. 4a) has features of a ‘fuzzy complex’(Forman-Kay and Mittag, 2013; Fuxreiter, 2018) where MB0 adopts multiple conformations in the bound state, in full agreement with the nature of the NMR intensity changes (Fig. 2). However, analysis of the orientation of MB0 in the 25,000 models with the lowest energy, shows a clear preference of direction of the peptide with the experimental-driven constraints compared to an analogous ensemble generated without constraints (Fig 4a; Supplementary Fig. 5d). Thus, despite the fuzzy nature of the ensemble including several clusters of opposite orientation, there is a clear preference for MB0 binding along the conserved patch of PAD from the center of α9 toward the center and C-terminus of α7 of PAD.

**Figure 4.**
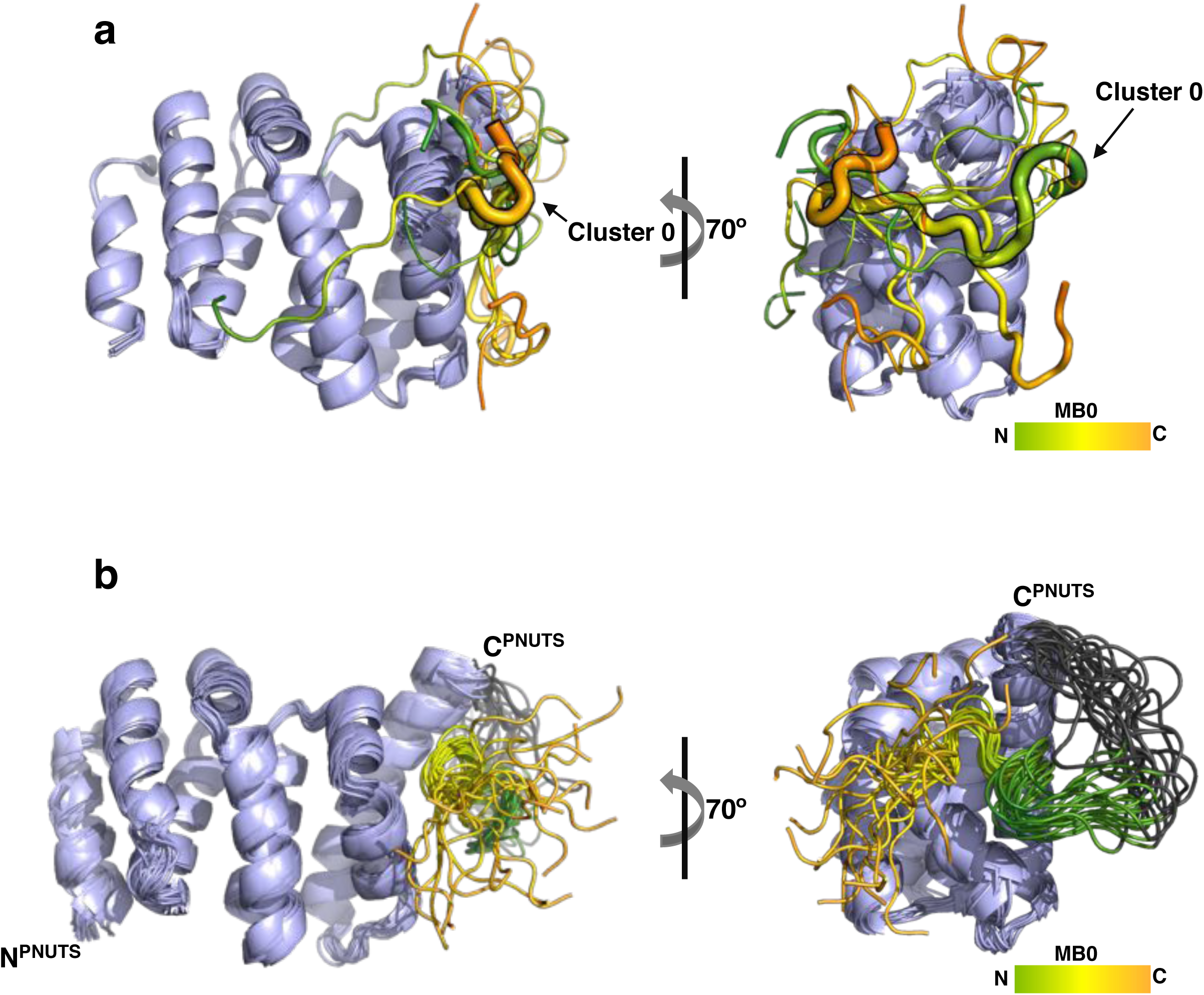
NMR guided model of PAD and MB0 interaction. (**a**) CSP-driven computational models of PAD binding to MB0 based on CSPs of MB0 and PAD obtained from the experiments depicted in Figure 2b and c and Figure 3c. Only the geometric cluster centre of 9 major MB0 conformation clusters are shown. Sizes of the peptides are proportional to what percentage of states belong to the cluster represented by the peptide. The dominant cluster 0 is marked with a bold outline and highlighted by an arrow. The PAD is colored light blue and MB0 is colored green to orange from N-termini to C-termini. (**b**) Experimental backbone trace of the NMR structure of the PAD-MB0 fusion protein (20 structures) with PAD, linker, and MYC_13-30_ colored light blue, grey, and green to orange, respectively. See also Figure S5.

### The solution structure of a PAD-MB0 fusion protein

To understand in greater detail how the PAD recognizes MB0, we engineered a fusion construct that could capture a predominant binding mode. This PAD-MB0 fusion protein (PAD-MB0-fusion) contains residues 1-148 of PNUTS and amino acids 13-30 of MYC with a flexible (GGGS)_2_ linker joining the two proteins. NMR analysis shows the PAD-MB0-fusion displays high quality, well dispersed NMR spectra (Supplementary Fig. 6a) enabling the full assignment of backbone and side-chain resonances. In comparing (^1^H-^15^N) HSQC spectra of apo PAD vs. PAD-MB0-fusion, the CSP patterns for PAD amides are similar to those observed when apo PAD is titrated with MB0 peptide (Supplementary Fig. 6b, c). For instance, several resonances in the ARM2 hydrophobic pocket that shift upon PAD binding to MB0 (e.g. M141, V117 and A114) follow a similar trajectory to their position in the spectrum of PAD-MB0-fusion; however these associated CSP amplitudes are significantly higher (Supplementary Fig. 6b). Differences in the perturbation patterns seen with PAD-MB0-fusion compared to those in the MB0 peptide binding assay are mainly localized to the PAD C-terminus close to the (GGGS)_2_ linker (Fig. 3c; Supplementary Fig. 6c, d). Taken together, these data suggest that the fusion construct favours MB0-bound states that are highly populated in the non-fused system, likely by increasing the local concentration of MB0 at the MYC-binding site of PNUTS.

Having validated the relevance of the PNUTS-MYC interaction within the fusion protein, we next performed a full, NOE-driven structure determination, including 43 experimental NOE restraints between the PAD and MB0 residues 13-30 (PDB ID: 7LQT, Fig. 4b, 5a, 5b; Table 1). As expected, MB0 binds on the conserved surface of ARM2 comprising helixes α7, α8, and α9, in good agreement with the NMR titration data (Fig. 3d, 4b). There are small differences between the structure of apo and PAD-MB0-fusion which superimpose with a backbone RMSD of 1.9±0.2Å (Supplementary Fig. 6d). The most important difference is confined to a small re-positioning of the C-terminal helix (α9) in the ARM2 motif, which we discuss in detail below.

**Figure 5.**
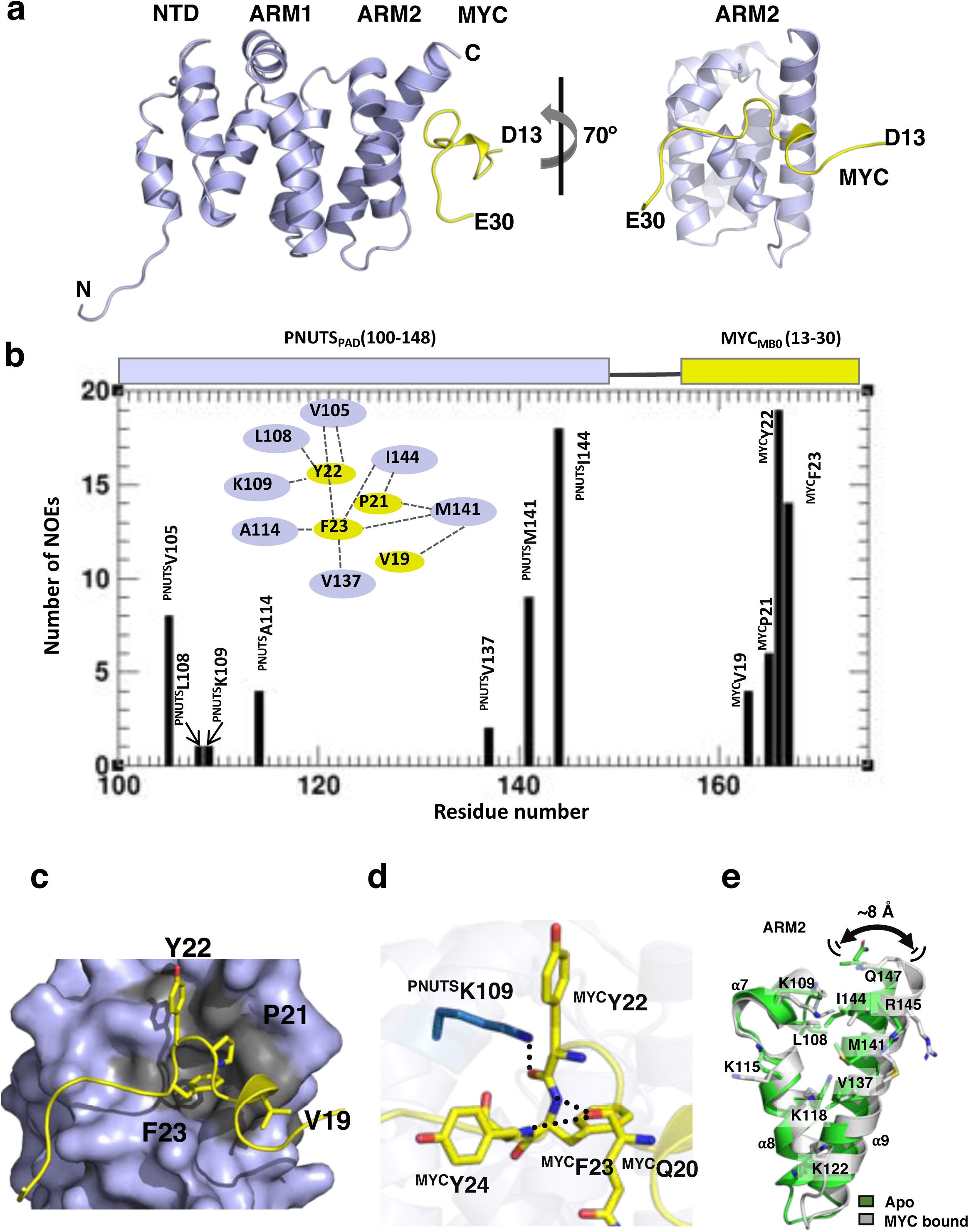
The specific interactions between PAD and MB0. (**a**) Cartoon representation of the solution state NMR structure of the PAD-MB0 complex, as derived from the PAD-MB0-fusion protein. The PAD is depicted in light blue and MB0 is =in yellow. (**b**) Histogram showing the number of assigned long-range NOEs between the PAD and MB0 residues, and vice-versa. The data was obtained from NOESY spectra of the PAD-MB0-fusion protein that is depicted above the diagram. Inset: Diagram of the distribution of inter-fragment NOEs - a dashed line connects MB0 and PAD residues with assigned NOEs. (**c**) Four residues within MB0 are crucial for the PAD-MB0 interaction within the fusion protein. MYC(13-30) is shown in cartoon (yellow) with the PAD ARM2 domain in surface representation (light blue). MYC residues V19, P21, Y22, and F23 are shown in stick format; the ring of F23 occupies the hydrophobic PNUTS pocket, which is formed by V105, A114, V137, W140, M141, and I144 (grey). The MYC fragment adopts a β-turn (residues 20-24) and a helical turn (residues 15-19). (**d**) Hydrogen bonds (dotted lines) and polar interactions (stick representations) are indicated on the PAD-MB0 interaction surface. PNUTS and MYC residues are colored in blue and yellow, respectively. One hydrogen bond is formed between PNUTS and MYC (PNUTS(K109)-MYC(Y22)). Moreover, two hydrogen bonds are formed within the MYC chain between MYC(Q20)-MYC(F23) and MYC(Q20)-MYC(Y22). (**e**) Cartoon representation of PNUTS-ARM2 in apo (green) and MYC-bound (grey) states. Helices α7 and α8 superimpose well, while helix α9 undergoes a conformational adjustment. Residues L108, K109, K115, K118, K122, V137, M141, I144, R145, and Q147 are shown in stick representation for reference. See also Figure S6.

There is excellent agreement between the solution structure of the PAD-MB0-fusion and the NMR-guided computational modeling of the interaction (Fig. 4a, b; Supplementary Fig. 5b, e). The position and direction of MB0 in all 20 structures of the fusion protein ensemble are consistent with the preferred binding ‘Cluster 0’ of the NMR-guided computational model (Supplementary Fig. 5a, d). Indeed, the inter-chain contact frequency of residues over the total number of computational models are also similar to the fusion protein, especially the largest cluster of docked models with a correlation R of 0.82 (Supplementary Fig. 5c).

### The specific interactions between PAD and MB0

Distinct hydrophobic and polar interactions are formed between the core hydrophobic motif of MB0-V_19_QPYFY_24_, and the conserved hydrophobic patch of PAD ARM2 (Fig. 5a). This interaction is defined by NOEs from four MB0 residues (V19, P21, Y22, F23) and PNUTS residues (V105, L108, K109, A114, V137, M141, I144) (Fig. 5b and 5c). This network of NOEs delineates a clear single mode of interaction between PNUTS and MYC that is very similar to that of Cluster 0 in the NMR-guided computational model (Fig. 5b and 5c). Notably, MYC(F23) sits in a hydrophobic pocket formed by L108, A114, V137, W140, M141, I144 and the acyl chain of K118 at the bottom of the C-terminal facing surface. A hydrogen bond between PNUTS(K109) and MYC(Y22), supported by NOEs, suggests additional binding specificity (Fig. 5b and 5d). Two hydrogen bonds within MB0 are also supported by NOEs between residues MYC(F23)-MYC(Y24) and MYC(F23)-MYC(Y22), forming a β-turn motif (Fig. 5d). The N- and C-termini of MB0 show no NOEs with any PAD residues, and are more disordered than the bound MYC-V_19_QPYFY_24_ in all 20 structures of the ensemble (Supplementary Fig. 6e).

Comparison of the MB0 interaction site of PAD in the absence (apo) and presence of MB0 reveals a subtle conformational adjustment of the three helices in the ARM2 motif upon MB0 binding in the fusion protein. Notably, this helix movement was also required in the NMR-guided computational models for MB0 binding to PAD, further supporting this observation (Supplementary Fig. 6g). In the MB0–bound state, helix α9 is slightly tilted away from helices α7 and α8 (Fig. 5e) translocating its C-terminus approximately 8 Å in the presence of MB0 compared to the apo form of PAD. As a result, key residues in □9, including binding pocket residues I144, V137, M141, are slightly distanced from helices α7 and α8 in the presence of MB0 (Supplementary Fig. 6f). Thus, PAD appears to accommodate MB0 binding by opening a binding pocket between helices 7-9 in ARM2.

### Mutation of key residues diminish PNUTS-MYC interaction *in vitro* and in human cells

To further validate the mode of interaction, we generated point mutations of residues in MB0 shown to be important to the PNUTS-MYC interaction (Fig. 3c and 5b) and then evaluated their binding to PAD in BLI assays (Fig. 6a; Supplementary Fig. 7). Alanine point mutants of the MB0 residues within the binding stretch V_19_QPYFY_24_ (Fig. 2) were generated within MYC(1-88) and evaluated for interaction with PAD by BLI. Interaction of wild-type MYC(1-88) with PAD has a binding affinity of K_D_ ∼3.5 µM (Fig 1b), whereas the MYC mutants MYC(1-88_P21A_), MYC(1-88_Y22A_), MYC(1-88_F23A_) and MYC(1-88_Y24A_) had reduced K_D_ values by a factor of 5 or more (Fig. 6a; Supplementary Fig. 7).

**Figure 6.**
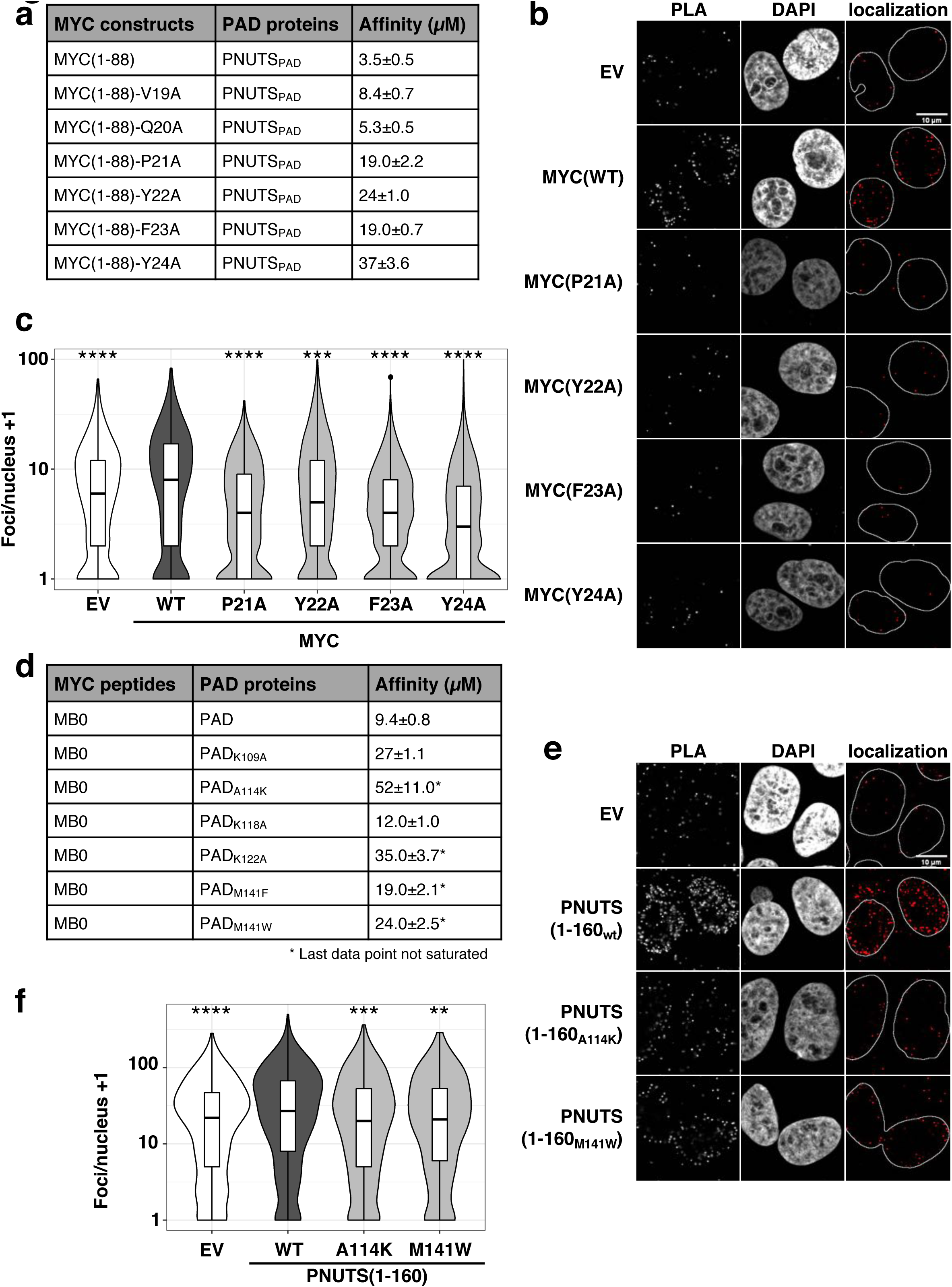
Mutation of key residues diminish PNUTS-MYC interaction *in vitro* and in human cells. **(a)** Key residues in the V_19_QPYFY_24_ stretch were individually mutated to alanine in the MYC(1-88) expression construct. The different MYC constructs were tested by BLI for binding to PAD and obtained affinities are displayed in the rightmost column. (**b**) Empty vector control (EV) or V5-tagged MYC constructs (Wild type (WT), P21A, Y22A, F23A or Y24A) were expressed in MCF10A cells and their proximity to endogenous PNUTS assessed by PLA. Representative images of magnified individual nuclei are shown. (**c**) Quantification of the in obtained images of (b) is shown as a violin plot. The plot is shown on a logarithmic scale and 1 was added to each measurement for visualization. Statistical significance was tested with the Kolmogorov-Smirnov test (***: p ≤ 0.001; ****: p ≤ 0.0001; n=3). (**d**) PAD residues that are important for MYC binding were substituted to amino acids that are able to disrupt the interaction while maintaining the folding of PAD. BLI was performed with MB0 peptide and measured affinities are displayed in the last column. (**e**) EV control, Flag-tagged PNUTS(1-160), or point mutants: PNUTS(1-160_A114K_), PNUTS(1-160_M141W_) were expressed in MCF10A cells expressing V5-MYC. PLA was performed using FLAG and V5 antibodies. Representative images of magnified individual nuclei are shown. (**f**) The foci formed per nucleus were quantified and are displayed on a logarithmic scale as foci/nucleus plus one. Statistical significance was tested using the Kolmogorov-Smirnov test (**: p ≤ 0.01; ***: p ≤ 0.001; ****: p ≤ 0.0001; n=3). See also Figures S7 to S9.

To assess whether these residues also contribute to the interaction of MYC and PNUTS in a cellular context, we evaluated the interaction using a proximity ligation assay (PLA). First, V5-tagged MYC and MYC point mutants were generated and stably expressed in the MCF10A breast epithelial cell line, which was one of many cell lines used to show functional relevance of the PP1:PNUTS-MYC axis(Dingar et al., 2018) and were chosen as they image well for PLA. Using V5 antibody, which was common to all constructs, we demonstrated similar levels of expression and nuclear localization of all mutants as well as wildtype MYC by immunoblot and immunofluorescence, respectively (Supplementary Fig. 8a, b). We then performed PLA to measure the interaction of MYC(WT), MYC(P21A), MYC(Y22A), MYC(F23A) and MYC(Y24A) with endogenous PNUTS (Fig. 6b). With this assay co-localization of two proteins of interest within approximately 40 nm is scored as a fluorescent focus. Several fields of view were assayed and the foci enumerated on a per nucleus basis. Indeed, each of the four MYC point mutants significantly reduced the number of fluorescent foci formed as compared to wildtype MYC (Fig. 6c). Overall, our mutational analysis confirmed that the residues on MYC, that we identified as key determinants of PNUTS-MYC interaction *in vitro,* are functionally important for this protein-protein interaction within cells.

We also evaluated the impact of single amino acid substitutions of PAD for their binding to MYC. Using FoldX (<u>foldx.crg.es</u>), amino acid substitutions were chosen to preferentially alter PNUTS-MYC interaction without disrupting protein structure. All six of the PAD point mutants (K109A, A114K, K118A, K122A, M141F and M141W) displayed weaker interaction with the MB0 peptide; in five of these, the binding affinity was reduced by more than 2-fold (Fig. 6d; Supplementary Fig. 7).

We similarly evaluated the degree to which the PAD can interact with MYC in human cell lines. To this end, we expressed a doxycycline-inducible, in-frame, fusion protein consisting of a Flag-tag, PNUTS amino acids 1 to 160 (PNUTS(1-160)) and three tandem SV40 nuclear localization signal (NLS) motifs. PNUTS(1-160) was used to achieve sufficient expression of the PAD in mammalian cells and the NLS ensured nuclear localization. Upon addition of doxycycline to the media, we observed the expected induction of PNUTS(1-160) as confirmed by immunoblot (Supplementary Fig. 9a) and nuclear localization determined by immunofluorescence (Supplementary Fig. 9b). To evaluate binding of PNUTS(1-160) to full-length MYC in human cells, we next performed PLA. Induction of PNUTS(1-160) dramatically increased the number of foci formed per nucleus compared to the EV control, giving us confidence that the PAD is also responsible for the interaction of MYC and PNUTS in human cells (Fig 6e). Evaluation of point mutants of key PAD residues important for MYC interaction, A114K and M141W, by PLA showed a significant reduction in foci formation as compared to wild-type PNUTS(1-160) (Fig. 6f). Taken together, these findings demonstrate that the interaction domains of MYC and PNUTS we mapped *in vitro* are also responsible for MYC:PNUTS interaction in human cells.

## Discussion

Distinguishing direct protein interactors of the MYC oncoprotein is essential not only to understand the mechanism of MYC function, but also to potentially unveil novel strategies to inhibit MYC binding to key protein partners and thus disable MYC as a potent cancer driver. We have previously shown the PP1:PNUTS phosphatase complex regulates MYC phosphorylation and degradation(Dingar et al., 2018), however the molecular basis of the interaction remained unclear. Here we show that PNUTS and MYC interact directly through the PAD and MB0 regions, respectively. Using NMR, the key residues of PAD-MB0 interaction were identified. The apo structure of the PAD was determined (PDB ID: 6VTI) and shown to consist of nine helices arranged as three helical bundles, with the two C-terminal bundles consisting of two ARM repeats; ARM1 and ARM2. Using NMR and computational modeling, we determined the molecular basis of the interaction not only by analyzing PAD and MB0 as well as PAD and MYC(1-88), but also a PAD-MB0-fusion protein (PDB ID: 7LQT). Our data support a model in which a core hydrophobic stretch in MB0 anchors onto the C-terminal facing surface of the PAD ARM2 motif primarily through hydrophobic interactions. Moreover, we show this interaction ensemble is highly dynamic but comprises a narrow range of specific, bound conformations. Validation of the interaction was achieved by demonstrating that point mutants of key interacting residues in either MYC or PNUTS disrupted the interaction both *in vitro* and *in vivo*.

The dynamic alterations in MYC in response to PAD binding, together with structural specifics of the bound state and *in vivo* effects of its inhibition, suggest a model in which MB0 binding to PAD could facilitate PP1:PNUTS access to phosphorylation sites within and adjacent to MBI for subsequent dephosphorylation (see Graphical Abstract). Our NMR analyses of MYC(1-88) showed that MB0 residues V19-Y24 comprise the primary anchor site of the PAD-MYC(1-88) interaction, with a weaker second touch-point within MBI (residues W50-L55). In contrast, MBI-neighbouring residues (P61 to V75) show dramatically sharper amide signals on PAD binding to MB0, suggesting a more rapidly interconverting ensemble state in this region compared to MYC(1-88) alone and a possible release of internal MYC interactions upon PAD binding, as we previously observed for tumour suppressor Bin1 binding to MBI(Andresen et al., 2012). In the context of the MYC-PNUTS interaction, the region of MYC that appears to be released upon PAD binding comprises critical serine/threonine residues essential for functional phosphorylation. This agrees with the role of the PP1:PNUTS-MYC axis in the regulation of MYC phosphorylation, activity and stability(Dingar et al., 2018). Such an anchor-release mechanism of PAD-MB0 binding in the PNUTS-MYC complex resembles that of Pin1 binding to MYC, where its anchoring to MB0 enhances cis-trans isomerisation c-terminal of MBI, thereby enhancing phosphorylation of S62(Helander et al., 2015). However, while Pin1 is a general cis-trans isomerase acting on many targets, PNUTS binding to MYC holds a specific key regulatory role of MYC function and degradation. In agreement, the specificity of the binding motif identified here is verified by single structure-based point-mutations that entirely abort the MYC-PNUTS interaction *in vitro* and *in vivo*.

Our analyses revealed that the structure of the PAD is also affected by the PAD-MB0 interaction. Specifically, in the MB0-bound state, the α9 helix of PNUTS ARM2 is tilted away from α8 compared to apo PAD. Both in-silico modeling and direct NMR studies suggest that this conformational change supports the formation of the PAD binding pocket required to accommodate the buried MYC(F23) residue. The PNUTS-MYC interaction may be further regulated by key PNUTS residues (I144, V137, M141) that could contribute to a more closed or open configuration of the ARM2 domain binding patch. Reciprocal structural changes in PNUTS that widen the binding groove upon MYC interaction suggest a mutual conformational adaptation, but whether this resembles induced fit or conformational selection models requires further investigation. Thus, our analysis reveals key structural and dynamic features of the PNUTS-MYC interaction which may jointly contribute to the regulation of the phosphorylation state of MYC.

PNUTS is one of several PP1 phosphatase regulatory subunits that determines substrate specificity. In particular, PNUTS directs PP1 dephosphorylation activity to protein substrates localized in the nucleus, including TOX high-mobility group box family member 4 (aka TOX4, LCP1)(Lee et al., 2009), phosphatase and tensin homolog (PTEN)(Kavela et al., 2013), telomeric repeat-binding factor 2(Kim et al., 2009), RNA polymerase II(Booth et al., 2000), retinoblastoma protein(De Leon et al., 2008), and WDR82(Landsverk et al., 2020). Of these only TOX4 and PTEN have been shown to bind to the PNUTS N-terminal region, which includes the PAD revealed in this work. The calponin-homology domain of TOX4 interacts with PNUTS residues 1-263(Lee et al., 2009), and the C2 tensin-type domain of PTEN has been reported to interact with the first 146 residues of PNUTS(Kavela et al., 2013). Here we show how the intrinsically disordered MYC(1-88) interacts with the PAD in a fuzzy complex, and identify and characterise a narrow ensemble of bound states by increasing the local concentration of MYC close to the binding site on PNUTS by means of a PAD-MB0-fusion protein. Taken together, these results suggest that the PAD recognizes and distinguishes specific PP1 nuclear substrates by a wide range of binding mechanisms, and thereby acts as a landing pad for protein interactions regulating the activity of the PP1:PNUTS complex.

Targeting MYC activity by disrupting MYC-protein interactions is an area of intense interest as inhibiting MYC directly has not been fruitful to date. As the bHLHLZ interaction with MAX was the first binding partner of MYC(Blackwood and Eisenman, 1991), several groups endeavour to exploit this partnership for the development of inhibitors(Carabet et al., 2018; Castell et al., 2018; Dang et al., 2017; Han et al., 2019; Lu et al., 2020). More recently, the focus has been on targeting interactors of the unique MYC Box regions. The interaction of MBIV with WDR5 induces a specific subset of genes whose products regulate protein synthesis, and inhibitors to block this interaction are under development(Chacon Simon et al., 2020; Thomas et al., 2019). Moreover, recent insight into the structural basis of MBII-TRRAP interaction has unveiled potential strategies to target this key interaction(Feris et al., 2019). Despite decades of MYC research, MB0 has only recently been recognized as a highly conserved MYC box that is functionally important for MYC oncogenic activity(Helander et al., 2015; Kalkat et al., 2018). Our finding here that PNUTS directly interacts with MB0 further reinforces the functional importance of this PP1:PNUTS-MYC regulatory axis, which we had previously shown controls MYC activity and stability(Dingar et al., 2018). Indeed, PNUTS was recently shown by an independent group to regulate N-MYC stability(Tee et al., 2020), further emphasizing the critical nature of the PNUTS-MYC interaction to MYC family activity. This is consistent with MB0 being conserved amongst MYC family members, thereby enabling PP1:PNUTS to regulate the rapid turnover of these oncoproteins. Importantly, MYC and PP1:PNUTS are often overexpressed in human cancers (Dingar et al., 2018; Marx et al., 2020) further supporting the concept that disrupting PNUTS-MYC by developing inhibitors targeting the PNUTS pocket in which MYC binds, may be possible, and have therapeutic utility. Such an inhibitor would not be generally cytotoxic as it would not target PNUTS *per se*, but would disrupt the PAD-MB0 interaction leading to MYC hyperphosphorylation and degradation. MYC-driven and/or - dependent cancers would be particularly vulnerable to the loss of MYC and undergo cell death. Given that MYC is dysregulated in the majority of human cancers, this strategy to decrease MYC activity makes the MB0-binding pocket of PNUTS an attractive drug target.

## ACKNOWLEDGEMENTS

This work was supported by funding from Canadian Institutes of Health Research (FRN156167 to L.Z.P.; FDN154328 to C.H.A.; FDN143312 to D.W.A.); Swedish Cancer Society and Swedish Childhood Cancer Fund (M.S.); Swedish Research Council (2018-04390 to M.S., 2016-05369 to B.W.); and the Princess Margaret Cancer Centre, The Princess Margaret Cancer Foundation, and Ontario Ministry of Health. The Structural Genomics Consortium is a registered charity (no: 1097737) that receives funds from; AbbVie, Bayer AG, Boehringer Ingelheim, Canada Foundation for Innovation, Eshelman Institute for Innovation, Genentech, Genome Canada through Ontario Genomics Institute [OGI-196], EU/EFPIA/OICR/McGill/KTH/Diamond Innovative Medicines Initiative 2 Joint Undertaking [EUbOPEN grant 875510], Janssen, Merck KGaA (aka EMD in Canada and US), Pfizer, Takeda and Wellcome [106169/ZZ14/Z]. The computations were performed on resources provided by the Swedish National Infrastructure for Computing (SNIC) at the National Supercomputer Centre (NSC) in Linköping. LZP and DWA hold Tier 1 Canada Research Chairs in Molecular Oncology and Membrane Biogenesis, respectively.

## Author contributions

Y.W., L.Z.P. conceived the project. Y.W. performed the bulk of protein purifications. V. M. cloned and purified MYC(1-88) for NMR. Y. W. performed BLI affinity measurements with the help from T.K.. Y.W., S.H., S.D., A.L., and A.A. performed the NMR studies of the apo PAD and PAD-MB0-fusion. A.L., S.H. determined the NMR structures of apo PAD and PAD-MB0-fusion protein. A.A., I. J.-Å., M.S., B.W. performed and evaluated the NMR studies of MYC peptides with PAD. I. J.-Å., A. A. and B.W. carried out the NMR-guided structural modeling. C.R., A.T. performed all experiments in human cell lines. D.W.A., B.W., M.S., C.H.A., L.Z.P. supervised the work. Y.W., A.A., C.R., I. J.-Å., S.H., D.W.A., B.W., M.S., C.H.A., L.Z.P. wrote the manuscript. All authors discussed the results and contributed to the final manuscript.

## STAR Methods

### Contact for Reagents and Resource Sharing

Further information and requests for resources and reagents should be directed to and will be fulfilled by the Lead Contact, Linda Z. Penn.

### Protein expression and purification

Several constructs within the 1-186 region of PNUTS were evaluated for expression by cloning into the pET28-MHL vector (RRID:Addgene_26096) containing an N-terminal His6 tag and a TEV cleavage site. PNUTS(1-148) was the shortest construct successfully expressed and stably purified in *E.coli* BL21 (DE3) cells. These cells were lysed by sonication with a Misonix S-3000 sonicator for a total processing time of 10 minutes with cycles of 5s sonication and 7s rest in 1X PBS buffer (PBS415, Bioshop). PNUTS was purified by Ni-NTA agarose (QIAGEN) washed with 1X PBS buffer with 5% glycerol and 2mM β-ME followed by gel filtration on Superdex 75/300 column (GE Healthcare), equilibrated with the NMR buffer containing 20 mM HEPES, pH 6.9, 200 mM NaCl, 2 mM DTT, 5% glycerol. Unlabeled proteins were grown in Terrific Broth medium and induced with 0.2 mM IPTG at 16 °C overnight. Double (^13^C-^15^N)-labeled PNUTS for NMR studies was grown in minimal M9 media supplemented with ^15^NH_4_Cl and ^13^C D-glucose.

For the PAD-MB0-fusion protein, cDNAs encoding PNUTS(1-148), a (GGGS)_2_ linker and MYC(13-30) were cloned in frame and inserted into pET15-Trx-MHL (modified by SGC based on pET15-MHL (RRID:Addgene_26096)), containing an N-terminal His_6_-tagged thioredoxin(Savitsky et al., 2010) and a TEV cleavage site. Thioredoxin was cleaved using TEV protease in dialysis buffer (1X PBS, 5% glycerol, 2mM β-ME) at 4°C overnight, after which the fusion protein was purified as described above.

MYC(1-88) used for NMR experiments was expressed from a pNH-TrxT vector (RRID:Addgene_26106) as a His_6_-tagged TEV cleavable fusion protein with Thioredoxin. Transformed BL21(DE3) cells (ROS-2, pRAR3 plasmid) were grown in LB medium and induced by 0.5 mM IPTG at 37° for 3 h. When OD_600_ was reached a level of 0.6, the cells were harvested by centrifugation, resuspended in lysis buffer (50 mM NaH_2_PO_4_, 10 mM Tris–HCl, 300 mM NaCl, 10 % glycerol and 20 mM β-ME at pH 8.0), and sonicated. The supernatant containing the protein was purified under native conditions with Ni-NTA (Invitrogen), cleaved with TEV, and finally purified using reverse IMAC and gel filtration (Superdex 75, Cytiva), dialysed into NMR buffer (20 mM HEPES, pH 7.0, 100 mM NaCl, 5 mM DTT, 5% glycerol) and concentrated. For preparing the (^13^C-^15^N)-labeled MYC(1-88), cells were grown in minimal M9 media supplemented with ^15^NH_4_Cl and ^13^C D-glucose and purified as mentioned above. To the final NMR sample 1 mM TCEP, 100 μM NaN_3_ and 10% D_2_O was added.

Biotinylated MYC(1-88) was expressed from a pET15-TrxT-MHL vector (RRID:Addgene_26106) containing an AVI tag following the MYC(1-88) cDNA. The protein was expressed in the BirA-transformed Competent BL21 (DE3)-BirA strain (designed by SGC), that co-expresses BirA protein ligases after IPTG induction. Protein was purified as described above with an extra wash step with wash buffer (1X PBS buffer with 5% glycerol and 2mM β-ME) including 1mM D-biotin. Mutations were introduced with the QuikChange II XL Site-Directed Mutagenesis Kit (Stratagene) and confirmed by DNA sequencing. Mutants were expressed and purified as the wild-type constructs above.

### Bio-Layer interferometry interaction measurements

Bio-Layer Interferometry (BLI) assays were performed using an Octet RED384 instrument (ForteBio, USA). C-terminally biotinylated MYC constructs or N-terminally biotinylated peptides were immobilized onto SA biosensors (ForteBio, USA) and dipped into serial dilutions of PAD (WT or mutant) in 384-well tilted-bottom microplates (ForteBio, USA). Assays were performed at room temperature in 1X commercial HBS-EP buffer (GE Healthcare, 10 mM HEPES pH 7.4, 150 mM NaCl, 3 mM EDTA, 0.005% Surfactant P20) supplemented with 0.002 mg/ml BSA. K_d_ values were calculated from concentration-dependent steady-state response curves, using Octet software v10.0.

### NMR Spectroscopy

NMR spectra were acquired on Bruker Avance spectrometers (operating at 600 and 800 MHz) and a Varian INOVA spectrometer (operating at 600 MHz) equipped with cryogenic probes. All spectra were processed with NMRPipe(Delaglio et al., 1995) and analyzed with SPARKY(Lee et al., 2015). Reconstruction of non-uniformly sampled 3D spectra was with mddNMR(Kazimierczuk and Orekhov, 2015). For PAD and PAD-MB0-fusion structure determinations, conventional 3D triple resonance backbone, and double resonance NOESY and TOCSY spectra were collected at 30°C as described previously(Lemak et al., 2011). Proteins were buffered in 20 mM HEPES (pH 6.9, 200 mM NaCl, 2 mM DTT, 5% glycerol, 7% D_2_O). Assignments were performed with aid of the software FMCGUI(Lemak et al., 2011). Assignments were 98.5% and 90.4% complete for backbone and sidechain resonances, respectively for PAD, and 96.4% and 90.2% complete for backbone and sidechain resonances, respectively for PAD-MB0-fusion. ϕ and ψ torsion angle restraints were derived from backbone chemical shifts using TALOS(Shen et al., 2009). Distance restraints were derived from cross peaks in NOESY spectra, and H-bond restraints, when applied, were for residues unambiguously determined to be in secondary structural elements based on NOE patterns, chemical shift assignments and backbone torsion angles. Automated NOE assignments and structure calculations were performed using CYANA 2.1(Guntert, 2004). The final 20 lowest energy structures were refined with CNSSOLVE by performing a short restrained molecular dynamics simulation in explicit solvent(Bollen et al., 2010); the resulting 20 structures comprise the NMR ensemble.

Binding studies, where (^13^C, ^15^N)-labeled PAD was titrated with unlabeled MYC(1-88) or biotinyl-MB0 peptide, with PAD to MYC molar ratios 1:1 and 1:2, and were performed at 30°C, buffered in 20 mM HEPES (pH 6.9, 200 mM NaCl, 2 mM DTT, 5% glycerol, 7% D_2_O). CSPs were calculated with ^15^N shifts weighted by a factor of 0.2 (for NH) and ^13^C shifts weighted by a factor of 0.17 (for CHα). Binding studies, where (^13^C, ^15^N)-labeled MYC(1-88) was titrated with unlabeled PAD, with PAD to MYC(1-88) molar ratios ranging from 0.03 to 1.9, and were performed at 15°C, buffered in 20 mM HEPES (pH 7.0, 100 mM NaCl, 5 mM DTT, 1 mM TCEP, 5% glycerol, 100 μM NaN_3_, 10% D_2_O) at PAD to MYC(1-88) molar ratios ranging from 0.03 to 1.9. Relative peak intensities (I/I_0_) were evaluated from HNCO spectra as described previously(Helander et al., 2015) and were normalized to five unaffected peaks to reduce the effect of dilution. NMR assignments for MYC(1-88) (untagged, natively expressed) used here were extrapolated from previous studies(Andresen et al., 2012; Helander et al., 2015) and confirmed by conventional 3D triple resonance backbone spectra..

### Molecular Docking of MB0 to the PAD

A model of the PAD-MB0 complex was constructed by docking the MB0 peptide guided by experimentally derived constraints using Rosetta FlexPepDock protocol(Raveh et al., 2011). The constraints were derived from chemical shift perturbations (CSP) and intensity differences from titration experiments, similarly to previous work modelling MB0 interacting with Pin1(Helander et al., 2015). For PAD, ^1^H-^15^N HSQC CSP were used from titration experiments with MYC(1-88) and biotinylated MB0, and additionally ^1^H-^13^C HSQC CSP was also used for MYC(1-88) titration. For MB0, ^1^H-^15^N HNCO CSP and intensity changes were used from titrating in PAD to MYC(1-88). The CSPs for the same nucleus and molecule were averaged, followed by the combination of all different CSPs to a single combined chemical shift perturbation, δ_i_ for each residue *i*, using a weighted average of the individual shifts with the weights based on the magnetogyric ratio of the nucleus; 1.000, 0.102, and 0.251 for ^1^H, ^15^N, and ^13^C(Schumann et al., 2007).

All combined chemical shift perturbations δ_i_ of residues differing by more than 2σ from 0 were included as constraints for the residues involved in binding. σ was calculated in an iterative manner as outlined in Schumann et al.(Schumann et al., 2007), by omitting combined shift perturbations, δ_i_,outside 3σ and recalculating until σ converged. The resulting shifts were normalized to a maximum signal of 1.0. For MB0, the HSQC intensity followed a similar process, but using σ from 1 instead. Relative intensities larger than 1.0 were set to 1.0. Resulting signals were reversely scaled from 1.0 to the lowest observed value. The actual significant combined shifts for MB0 were averaged with the significant signals from the peak intensity to form the final significant shifts for MB0. In preparation for usage as constraints, combined chemical shifts δ_i_ were reduced to a representative set of significant shifts by the iterative process as described above, and reweighted so the different chemical shift perturbation and intensity difference experiments contributed equally to the final constraints.

Pairwise ambiguous constraints were constructed between every exposed PAD residue (> 5% relative surface exposure) with significant combined shift to every MB0 residue with significant shift by pairwise addition of the signals. The constraints were imposed during the docking using a square well scoring function that positively favour the atom pair when within 5.0 Å, and linearly decreasing the constraint influence in the distance range 5 Å to 10 Å, while not penalizing distances further than 10 Å. This corresponds to the FADE constraint function type in Rosetta with a lower bound -5.0, upper bound 10.0, and cubic splines of width 5.0. The overall constraint weight was set to contribute as much to the final scoring of the complex as all the unconstrained energy functions.

The PAD structure was energy-minimized using the Rosetta relax protocol. 50,000 decoys of PAD:MB0 interactions were generated using Rosetta Flex-PepDock ab-initio protocol guided by the constraints. Each decoy was generated starting from an extended MB0 peptide superpositioned on one of the constraint-pairs in a random orientation on the receptor surface. Rosetta FlexPepDock utilizes Monte-Carlo sampling to minimize the ‘REF2015’ energy function(Raveh et al., 2010), while sampling the translational, rotational, and torsional degrees of freedom of both MB0 and PAD. The translational and rotational degrees of freedom are sampled by randomly moving and rotating MB0 with respect to PAD. The backbone torsions of MB0 are sampled using fragment insertions, where the fragments are 3-, 5-, or 9-residue backbone dihedrals with local sequence similarity to the MB0. After each structural change, the side chains are rebuilt using Dunbrack’s backbone-dependent rotamer library(Shapovalov and Dunbrack, 2011) followed by energy minimization before the trial energy is calculated. Following the standard Metropolis acceptance criterion, the structural change is accepted with a probability related to the difference in energy before and after the change.

To properly represent the plausible conformations of the MB0 peptide, a subset of representative decoys which best describe the observed CSP were selected. First, the 25,000 models (median filter) with best resulting energy scores were clustered by backbone position, with cluster radii ranging from 1.0 to 4.0 Å in increments of 0.25, ignoring the flexible loops on the receptor when superpositioning and calculating RMSD. Then, each model in a cluster was described as a binary vector denoting for each residue the involvement in inter-chain interaction, defined as an inter-chain distance between any pair of non-hydrogen atoms ≤ 4.5 Å. Each cluster was represented by the sum of all member vectors. The Lasso algorithm as implemented in scikit-learn(Pedregosa, 2011), a linear model with L1 regularizer, was used to find the combination of clusters that had the best square-sum fit to the experimentally derived constraints. This was repeated for all cluster radii from above and the cluster radius 1.75 Å obtained the best fit to the constraints, with 9 clusters together describing the constraints with R^2^ 0.71 (correlation R of 0.84).

### Cell lines

Cell lines were grown as previously described(Wei et al., 2019). To evaluate the effect of MYC point mutations on the interaction with endogenous PNUTS, MCF10A cells were transduced with pLenti neo CMV V5-MYC or the respective mutants (P21A, Y22A, F23A, Y24A) and selected in 750 µg/mL Neomycin (Gibco). To test the interaction of PNUTS(1-160) wildtype or point mutants with MYC in cells, the MCF10A cell line was sequentially transduced with: pLenti neo CMV V5-MYC; pLenti blast CMV rtTA and pLenti hygro CMV/tight EV; and Flag-PNUTS(1-160)-3xNLS wild type or point mutants (A114K, M141W). After each transduction, cells were selected with the respective antibiotic (Neomycin/Geneticin: 750 µg/mL (Gibco), Blasticidin: 5 µg/mL (BioShop), Hygromycin B: 100 µg/mL (BioShop)). All cell lines were validated by STR profiling and tested regularly for mycoplasma using the MycoAlert Kit (Lonza) and remained negative throughout.

### Protein extraction and immunoblotting

The protein expression of both MYC and PNUTS(1-160) and point mutants was assessed after selection and for each replicate of PLA. To this end, cells were seeded in culture media and expression was induced with 0.5 µg/mL doxycycline (Sigma-Aldrich) for 24 h prior to protein harvest, where applicable. For PNUTS(1-160) constructs, cells were treated with 10 µM MG132 (Calbiochem) 4h prior to harvest. Cells were washed twice with 1x PBS (Wisent) and lysed on the plate using SDS lysis buffer (160 mM Tris-HCl pH 8.8, 2% SDS). Protein lysate was quantified using the Pierce 660 nm kit with added Ionic Detergent Compatibility Reagent (Pierce) and 10-15 µg of total cell lysate were run on a 10% SDS-PAGE. Proteins were wet transferred onto 0.2 µm Nitrocellulose membranes (Perkin Elmer) at 100 V for 1 h in chilled transfer buffer. Membranes were blocked with 5% skim milk in 0.1% PBS-T and incubated overnight with mouse anti-Flag (B3111, Sigma-Aldrich, 1:500), mouse anti-V5 (ab27671, Abcam, 1:1000), or rabbit anti-Actin (Sigma-Aldrich, 1:3000) in 5% skim milk in PBS-T. Detection was performed using fluorescently labeled secondary antibodies against rabbit and mouse (LI-COR) on the LI-COR Odyssey imaging system.

### Proximity Ligation Assay

The expression of the PNUTS constructs was induced through the addition of 0.5 µg/mL doxycycline (Sigma-Aldrich) for 24 h and 10 µM MG132 (Calbiochem) 4 h prior to fixation. For all PLAs with PNUTS(1-160) and its mutant, cells were fixed 10 minutes in 4% paraformaldehyde (Sigma-Aldrich) in PBS at pH 7.2. PLAs for MYC and MYC point mutants were fixed with 2% paraformaldehyde (Sigma-Aldrich) in PBS at pH 7.2 for 20 minutes.

Rabbit anti-MYC (Millipore, 06-340), mouse anti-Flag (Sigma, B3111), rabbit anti-V5 (Millipore, AB3792) and mouse anti-PNUTS (BD Bioscience, 611060) antibodies were optimized and used at a 1:200 (MYC, Flag), 1:500 (V5) or 1:50 (PNUTS) dilution, respectively. The PLA was performed as previously described(Dingar et al., 2015). The fluorescent foci were quantified using the Blobfinder software (Carolina Wählby & Amin Allalou, CBA, Uppsala University), with a minimum of 200 nuclei per biological replicate quantified. The data was plotted using R (version 4.0.2) using the function *ggviolin* from the package ggpubr (version 0.4.0). Statistical analysis was performed using the *ks.test* command in R.

## ACCESSION NUMBERS

Atomic coordinates and structure factors for the reported NMR solution structures have been deposited with the Protein Data Bank under accession number 6VTI and 7LQT.

## SUPPLEMENTAL INFORMATION

Supplemental Information includes nine figures and can be found with this article online at [link to be added].

---

**SI Figure S1.**
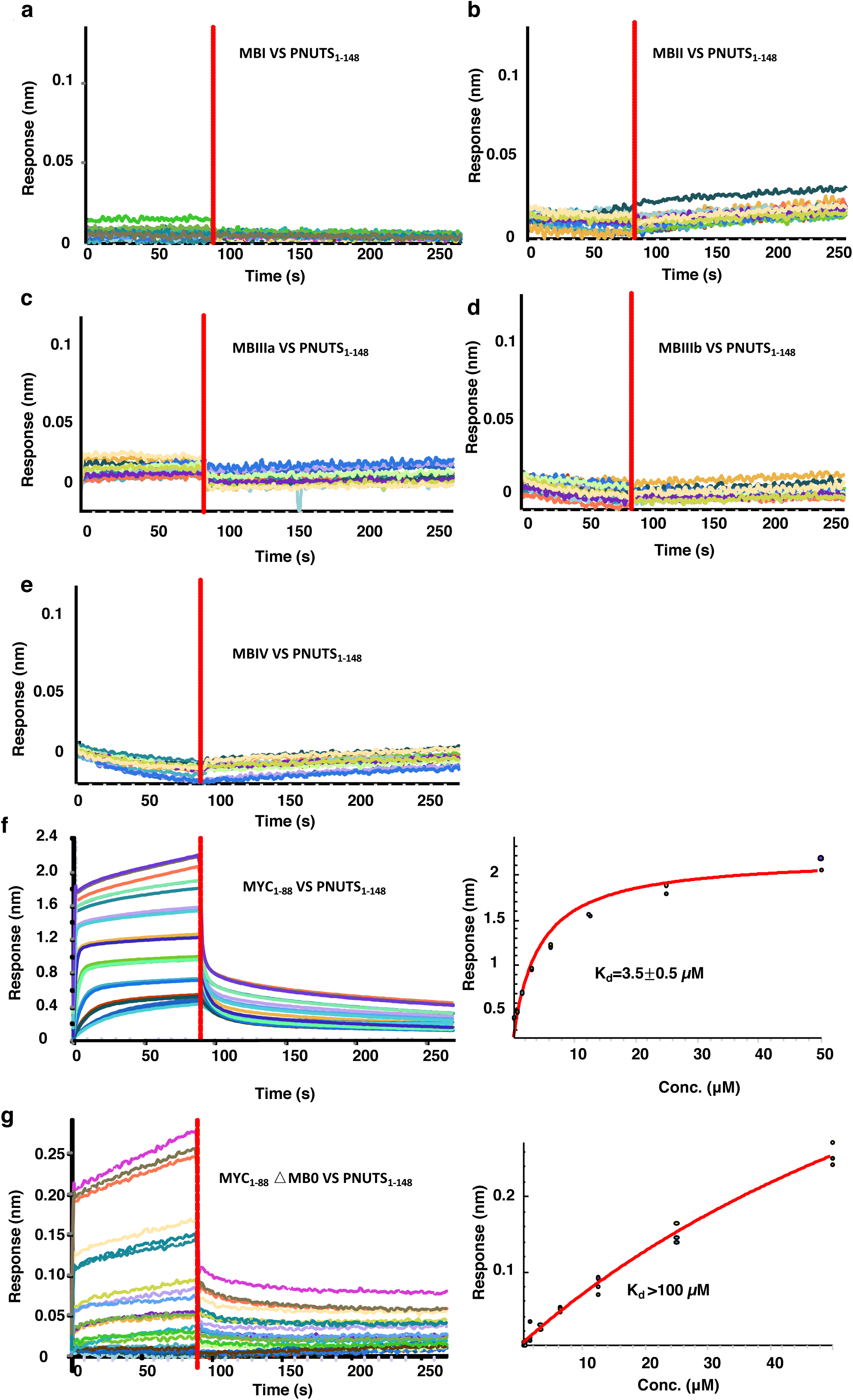
BLI sensorgrams for the interaction of PNUTS(1-148) with sensor-immobilized (a) MBI (Biotin-GKAPSEDIWKKFELLPTPPLSP), (b) MBII (Biotin-IIIQDCMWSGFSAAAK), (c) MBIIIa (Biotin-CIDPSVVFPYPL), (d) MBIIIb (Biotin-EEEIDVVSVEKR), and (e) MBIV (Biotin-THQHNYAAPPSTRKDYPAAKR), (f) MYC_1-88_ (left) and steady-state fit (right), (g) MYC_1-88ΔMB0_ (left) and steady-state fit (right).

**SI Figure S2.**
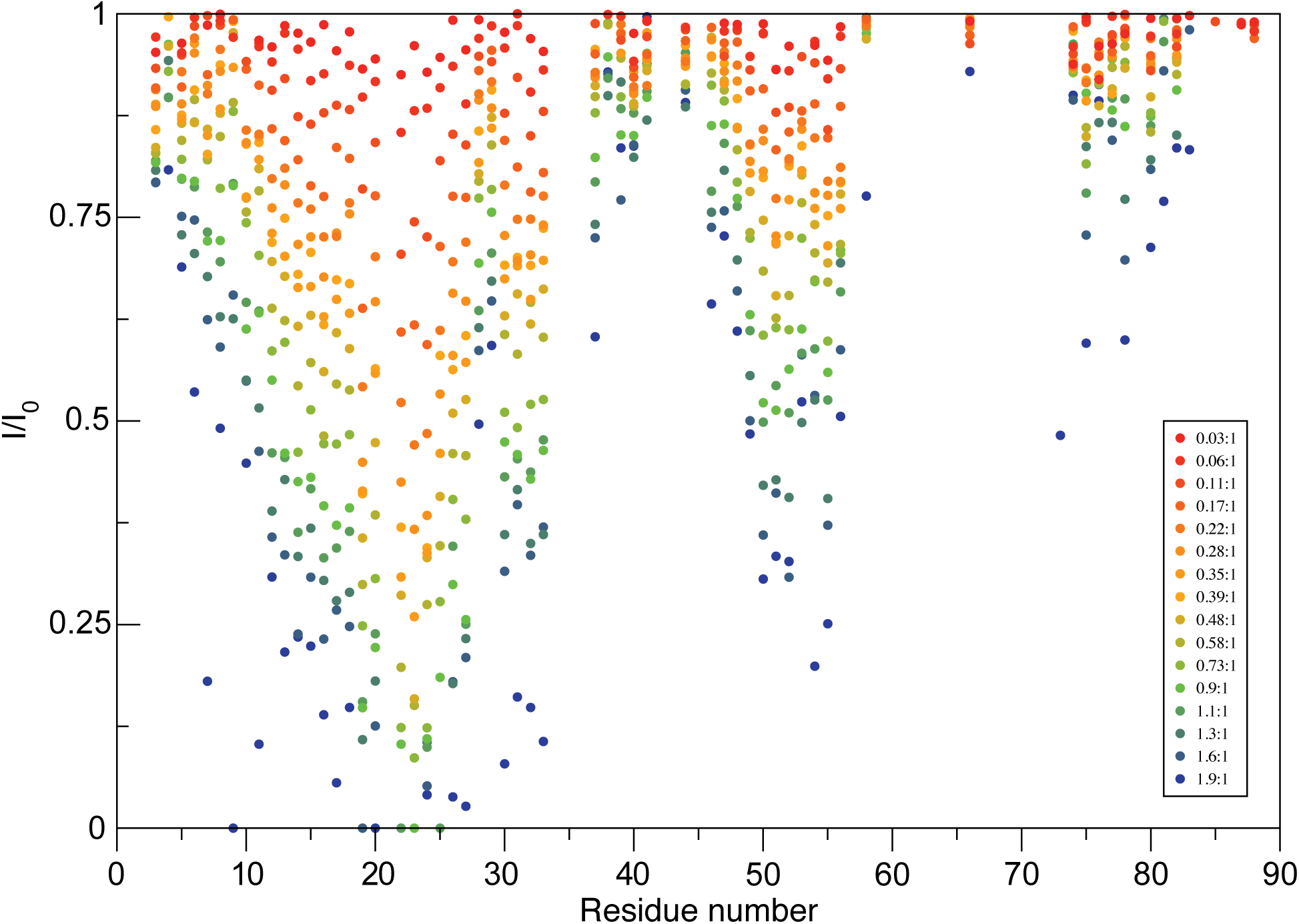
Changes in intensity of MYC(1-88) amide resonances (from HNCO spectra) upon addition of PAD (at MYC to PAD molar ratios ranging from 0.03 to 1.9.

**SI Figure S3.**
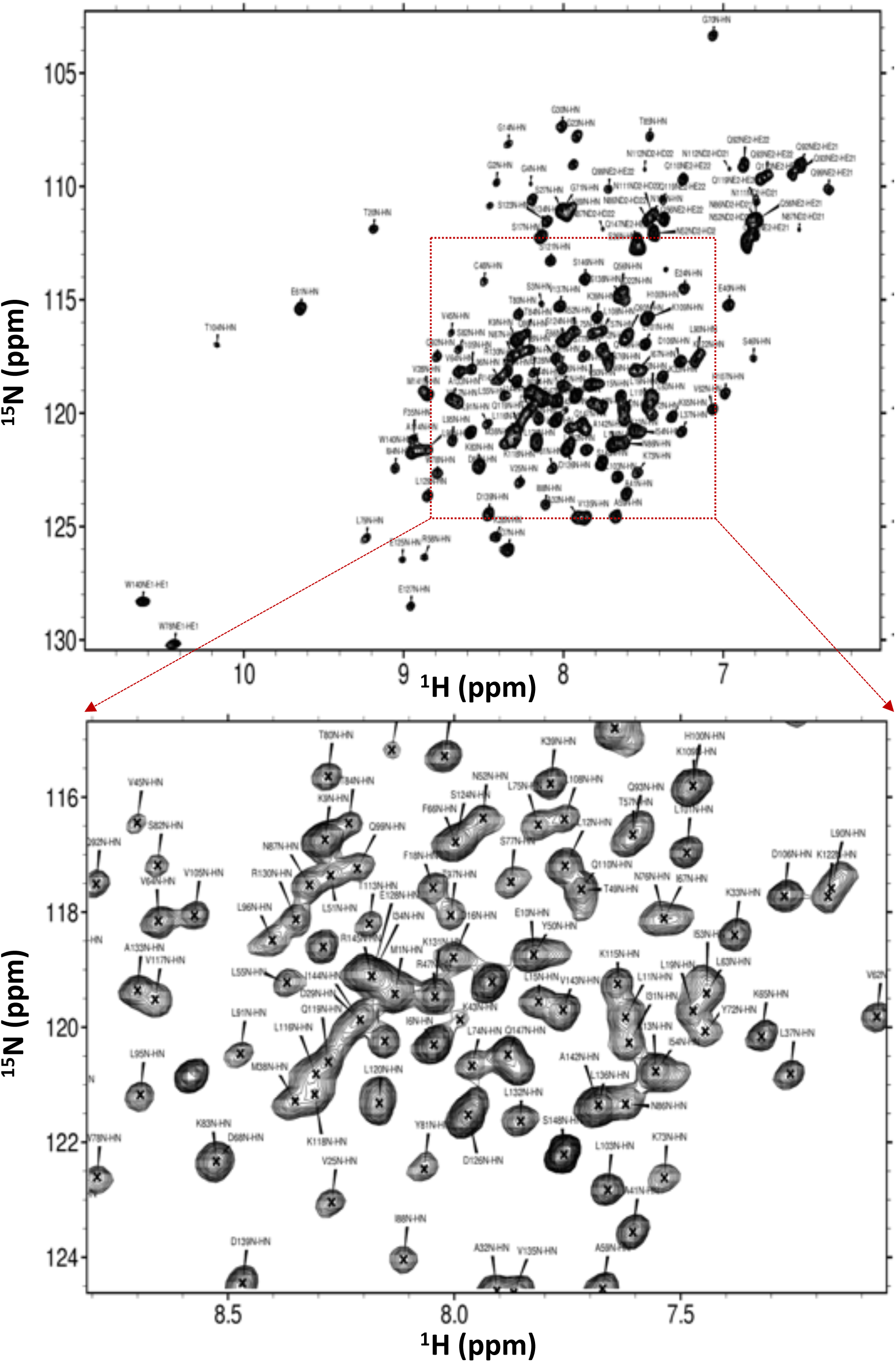
Full (top) and centre-zoomed (bottom) (^1^H-^15^N) HSQC spectrum of PAD (450uM, pH 6.9, 30°C).

**SI Figure S4.**
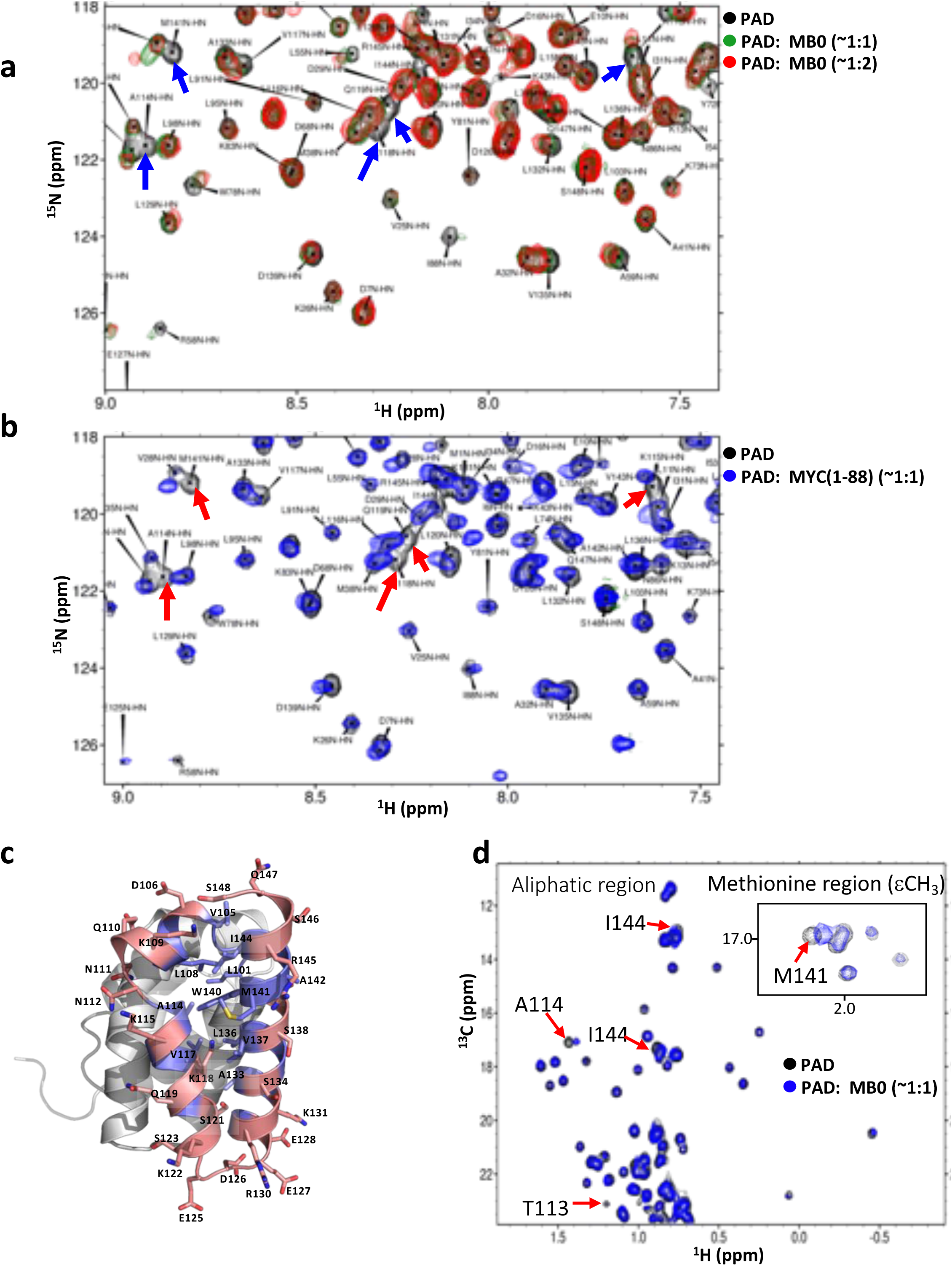
Identifying the PNUTS-MYC interface. (a) (^1^H-^15^N) HSQC overlay of PAD (apo – black) upon binding to biotinyl-MB0 peptide (MYC_16-33_) at PNUTS:MYC molar ratios of 1:1 (green) and 1:2 (red). Arrows point to residues with significant CSPs. (b) (^1^H-^15^N) HSQC overlay of PAD (apo – black) upon binding to MYC(1-88) at a molar ratio of 1:1 (blue). Arrows point to residues with significant CSPs. (c) Residues in/around the C-terminal facing surface are shown in sticks. Aromatic residues are displayed in light-blue, polar residues in salmon. (d) Methyl region of a CT (^1^H-^13^C) HSQC overlay of PAD (apo – black) upon binding to MYC(1-88) at a molar ratio of 1:1 (blue). Residues with observable CSPs are annotated.

**SI Figure S5.**
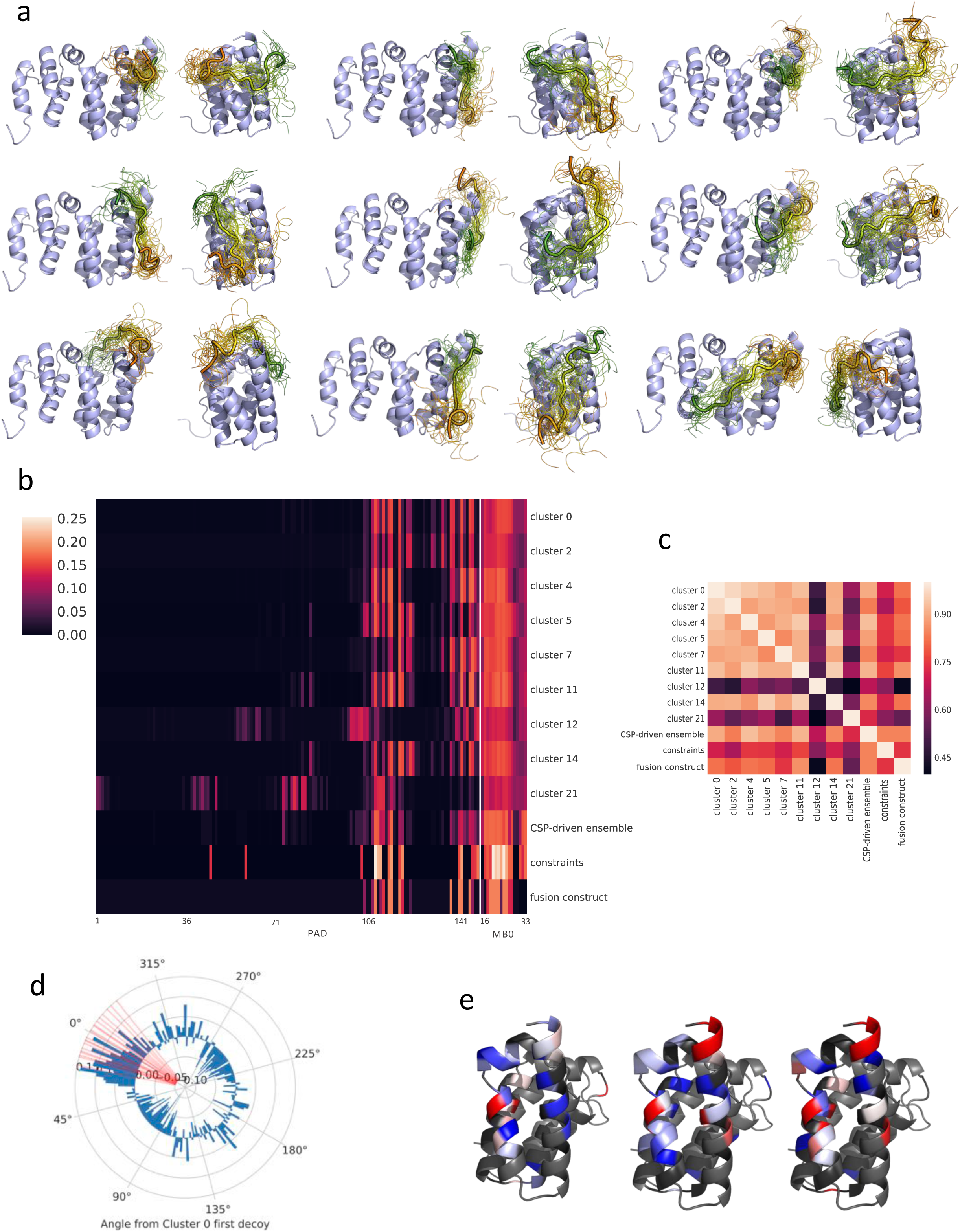
Modeling of PAD-MB0 complex. (a) 30 lowest-energy MB0 conformations for each major conformation cluster of the CSP-driven complex models, colored as in Figure 4a. Geometric cluster center in larger size with bold edges. From left to right, up to down, the clusters are: 0, 2, 4, 5, 7, 11, 12, 14, and 21. (b) Contact preference profiles for the 9 major MB0 conformation clusters of the CSP-driven complex models, as well as the constraints used in the CSP-driven modeling, and the contact preference profile for the 20 lowest energy models of the fusion protein construct. Color intensity indicates what fraction of the models in the cluster display an inter-chain contact involving each residue. (c) Correlations between the contact preference profiles of the major MB0 conformation clusters, the combined final ensemble of modeled conformations, the CSP-derived constraints, and the contact preference profile of the 20 lowest energy models of the PAD-MB0 fusion protein construct. (d) Histogram over preference of direction of binding of the MB0-PYFY motif implied by CSP-derived constraints. Calculated as histogram of direction of binding of the MB0-PYFY motif in the MYC-binding groove of PNUTS guided by CSP-derived constraints subtracted with the same histogram without CSP-derived constraints (only using Rosetta peptide docking energy function). The angles of high fraction indicate that including the CSP-derived constraints biases the model towards these binding angles, while a negative fraction indicates the model is biased away from these angles. Direction angle measured from geometric center of CSP-driven complex cluster 0. Direction of the MB0-PYFY motif in fusion protein construct model marked in red. (e) Differences in chemical shift perturbation direction visualized for the MB0 binding groove on PAD. From left to right: difference between PAD to MYC(1-88) and PAD to biotinylated MB0, difference between PAD to MYC(1-88) and PAD-MB0 fusion protein, and difference between PAD to biotinylated MB0 and PAD-MB0 fusion protein. A color ranging from white to red indicates a difference in chemical shift perturbation direction, with clear red indicating an angle of 30 degrees or more. Black coloring indicates the residue has significant chemical shift perturbation for one of the complexes, but not the other, while gray coloring indicates none of the complexes have significant chemical shift perturbations for that residue.

**SI Figure S6.**
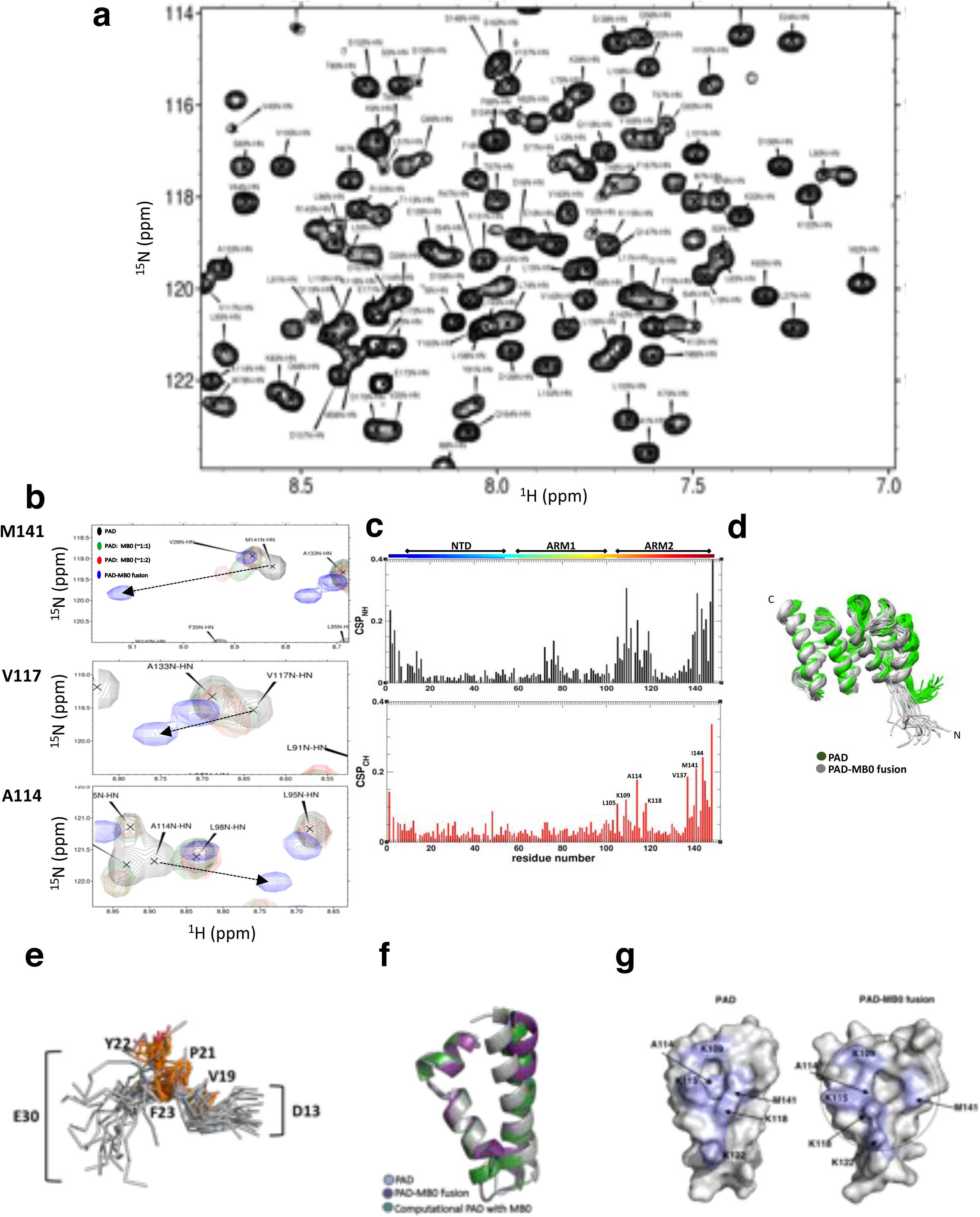
Complex structure of PNUTS-MYC. (a) Center-zoomed ^1^H-^15^N HSQC spectrum of PAD-MB0 fusion construct (450 µM, pH 6.9, 30°C). (b) Position of M141, V117 and A114 resonances in HSQC spectra of PAD (apo – black) upon titration with MB0 (at PNUTS to MYC molar ratios of 1:1 and 1:2) and in the PAD-MB0 construct. (c) Comparison of the chemical shifts of the residues of PAD measured in apo and fusion protein. Difference between backbone NH resonances is shown on the top panel. The maximum difference between aliphatic CH resonances calculated for atoms in a given residue is shown on the bottom panel. (d) Alignment of MB0-bound (gray) and apo PAD structures (alignment is based on residues 12-145). Differences can be seen at the N-terminus and in the ARM2 motif. The pairwise RMSD is 1.3 Å when both N-terminal and C-terminal helices are excluded and increases to 1.9 Å when they are included. (e) Conformations of key residues of MB0 involved in the PAD-MB0 fusion protein interaction in all 20 conformers. Four residues (V19, P21, Y22, and F23) involved in the hydrophobic interactions are shown in orange. (f) Comparison of the ARM2 in Apo, MB0 bound and computational states. (g) The exposed surface of the ARM2 pocket (dotted line) in apo (**left**) *vs.* MB0-bound PAD (**right**). The positions of PAD residues K109, A114, K115, K118, K122, and M141 in both structures are colored in light blue.

**SI Figure S7.**
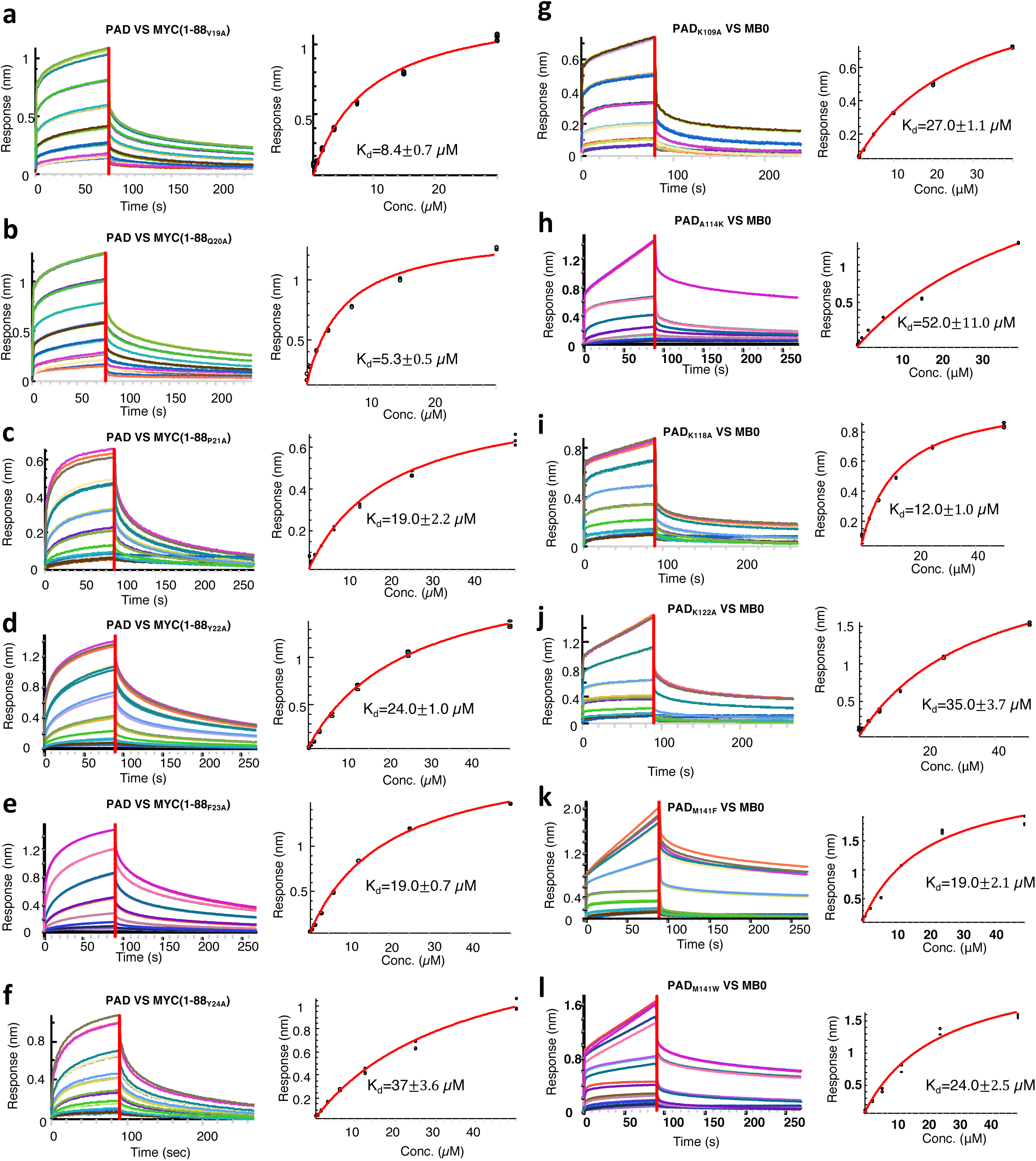
BLI sensorgrams and steady-state fits for the interaction of mutants of both PAD and MB0. (Left, a-f) Interactions between PAD and mutants of MYC(1-88). (Right, g-l) Interactions between MYC-MB0 and mutations of PAD.

**SI Figure S8.**
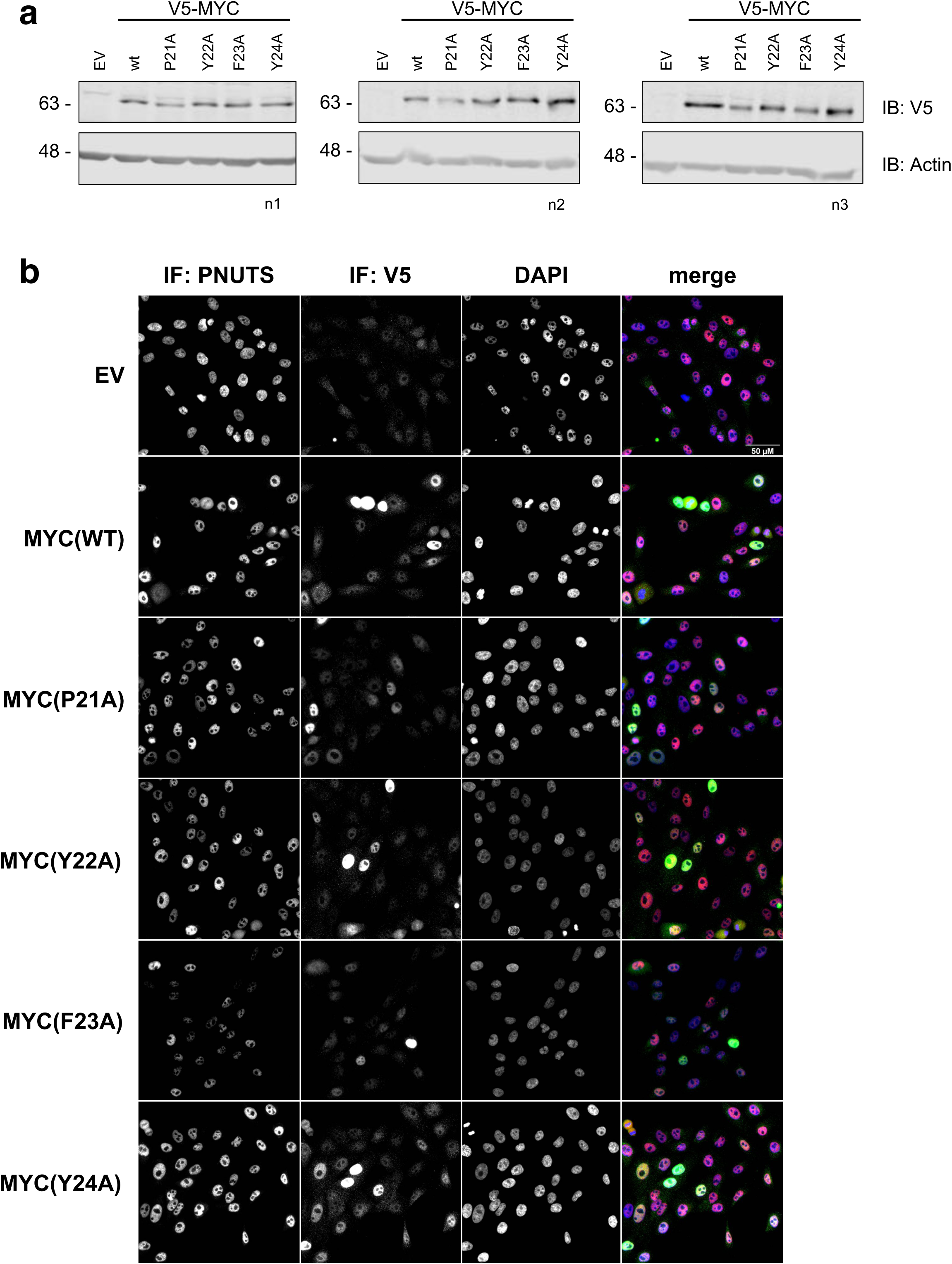
Expression of MYC point mutants in MCF10A cells. (a) Western Blot for protein expression of V5-tagged MYC and respective point mutants. Actin was used as a loading control. Protein lysates were harvested at the same time as PLA was performed. (b) Representative immunofluorescence images of cells stained with V5 (green) to detect V5-tagged MYC or respective point mutants and endogenous PNUTS (red) in MCF10A cells. Localization and expression are comparable between the EV (empty vector), WT (wild type) and mutant MYC cell lines.

**SI Figure S9.**
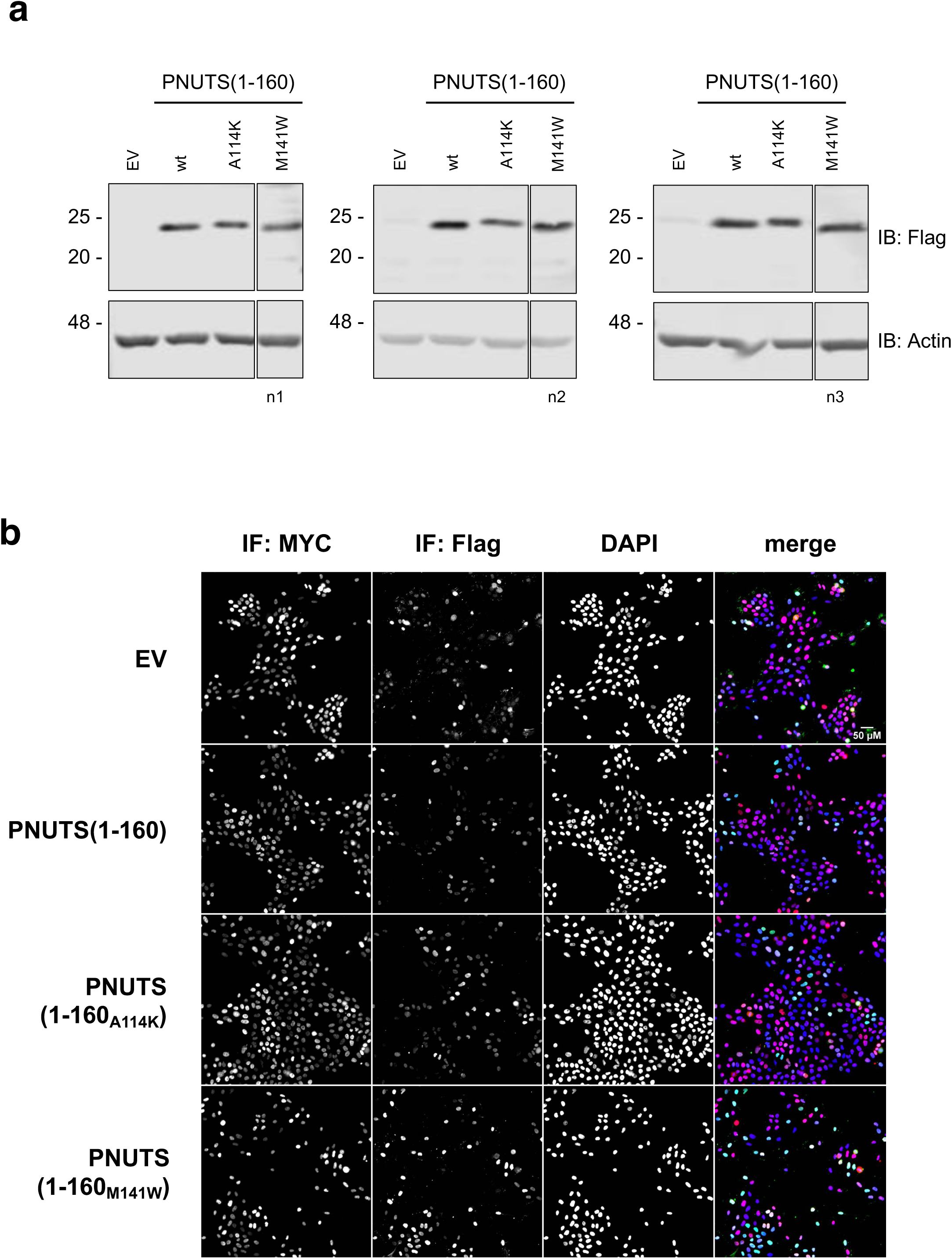
Expression of PNUTS(1-160) and point mutants in MCF10A cells. (a) Western Blot for protein expression of V5-tagged MYC and Flag-tagged PNUTS(1-160), PNUTS(1-160_A114K_) or PNUTS(1-160_M141W_). Protein lysates were harvested at the same time as PLA was performed. Actin was used as a loading control. (b) Representative immunofluorescence images taken as a control for the PLA between MYC (red) and Flag-PNUTS(1-160) as well as the A114K and M141W point mutants of Flag-PNUTS(1-160) (green). Cells were stained with DAPI to confirm the nuclear localization of the PNUTS(1-160) proteins. Overall, localization and expression are comparable between the WT and mutant PNUTS(1-160).

